# Heterogeneous aggregation of amyloid-β 42 from single-molecule spectroscopy

**DOI:** 10.1101/2020.09.10.290023

**Authors:** Fanjie Meng, Janghyun Yoo, Hoi Sung Chung

## Abstract

Protein aggregation is implicated as the cause of pathology in various diseases such as Alzheimer’s and Parkinson’s disease. Polymorphism in the structure of fibrils formed by aggregation suggests the existence of many different assembly pathways and therefore a heterogeneous ensemble of soluble oligomers. Characterization of this heterogeneity is the key to understanding the aggregation mechanism and toxicity of specific oligomers, but in practice it is extremely difficult because oligomers cannot be readily separated. Here, we investigate highly heterogeneous oligomerization and fibril formation of the 42-residue amyloid-β peptide (Aβ42). We developed and used new single-molecule fluorescence spectroscopic and fluorescence lifetime imaging methods, combined with deep learning for image analysis. We found that the concentration of oligomers, including dimers, is extremely low and that the dimer is conformationally diverse. Aggregation to form fibrils is also highly heterogeneous in terms of the number of strands in a fibril and the elongation speed and conformation of fibrils. This heterogeneity in all stages of aggregation explains diverse and sometimes irreproducible results of experimental studies of amyloid-β. Based on our observations and analysis, we propose a new model for aggregation of Aβ42.

## Introduction

Amyloid-β (Aβ) is a peptide fragment consisting of 39 – 43 amino acid residues, which is produced by successive proteolytic cleavages of the amyloid precursor protein (APP)^1^. Its aggregation to form fibrils that are found in brain tissue is one of the key characteristics of Alzheimer’s disease. Although fibrils are known to be toxic, a number of experimental studies also suggest that a subset of soluble oligomers, transiently appearing during the aggregation process, are more responsible for disease pathology than the fibril itself^2–6^. To better understand the pathogenic mechanism, therefore, it is important to characterize the entire aggregation process from the initial oligomerization to the formation and growth of fibrils.

Despite tremendous effort to understand aggregation of Aβ to form oligomers and fibrils, experimental results vary widely and there is no consensus on the model for these processes^7^. One of the difficulties in studying Aβ may result from its hydrophobicity because it is a part of the transmembrane domain of APP. Aβ interacts with many proteins either specifically or non-specifically as a monomer and various oligomeric forms^8^. A more fundamental reason for the difficulty may be the heterogeneity of the aggregation process. Aβ forms amyloid fibrils, long fibers with parallel (or anti-parallel) β-sheet structures (cross-β structure). Interestingly, there are variations in the fibril structure^9,10^ depending on various factors such as aggregation conditions. This polymorphism indicates that the entire aggregation process, including oligomerization, should be heterogeneous, which complicates biophysical and biochemical characterizations of Aβ. Due to the lack of quantitative experimental results, there is no comprehensive aggregation model that includes heterogeneity of aggregation pathways.

Single-molecule spectroscopy can be an effective tool to probe heterogeneity because molecular species can be observed one at a time, so individual oligomers can be detected without separation. This technique has the potential to identify toxic species. In this initial study, we combine single-molecule Förster resonance energy transfer (FRET) spectroscopy, fluorescence lifetime imaging (FLIM), and image analysis using deep learning (codes are available at https://github.com/hoisunglab/FNet) to interrogate several steps during the aggregation process of the 42-residue Aβ peptide (Aβ42), including dimerization, formation of stable oligomers, and fibril elongation.

## Results

### Dimerization

In our previous work using single-molecule FRET and MD simulation, we have shown that the monomer of Aβ42 is almost completely disordered with no structured regions that could possibly be a template for fibril formation^11^. The first step of oligomerization is most probably the formation of Aβ dimer. To selectively monitor the dimerization without interference from further oligomerization, we immobilized donor (Alexa 488)-labeled Aβ42 on a glass surface and incubated it with acceptor (Alexa 594)-labeled Aβ at a relatively low concentration (300 nM, Fig. 1a). (see Methods in the Supplementary Information and Supplementary Fig. 1 for the design of Aβ42 constructs). The experiment was finished before aggregation occurs.

**Fig. 1.**
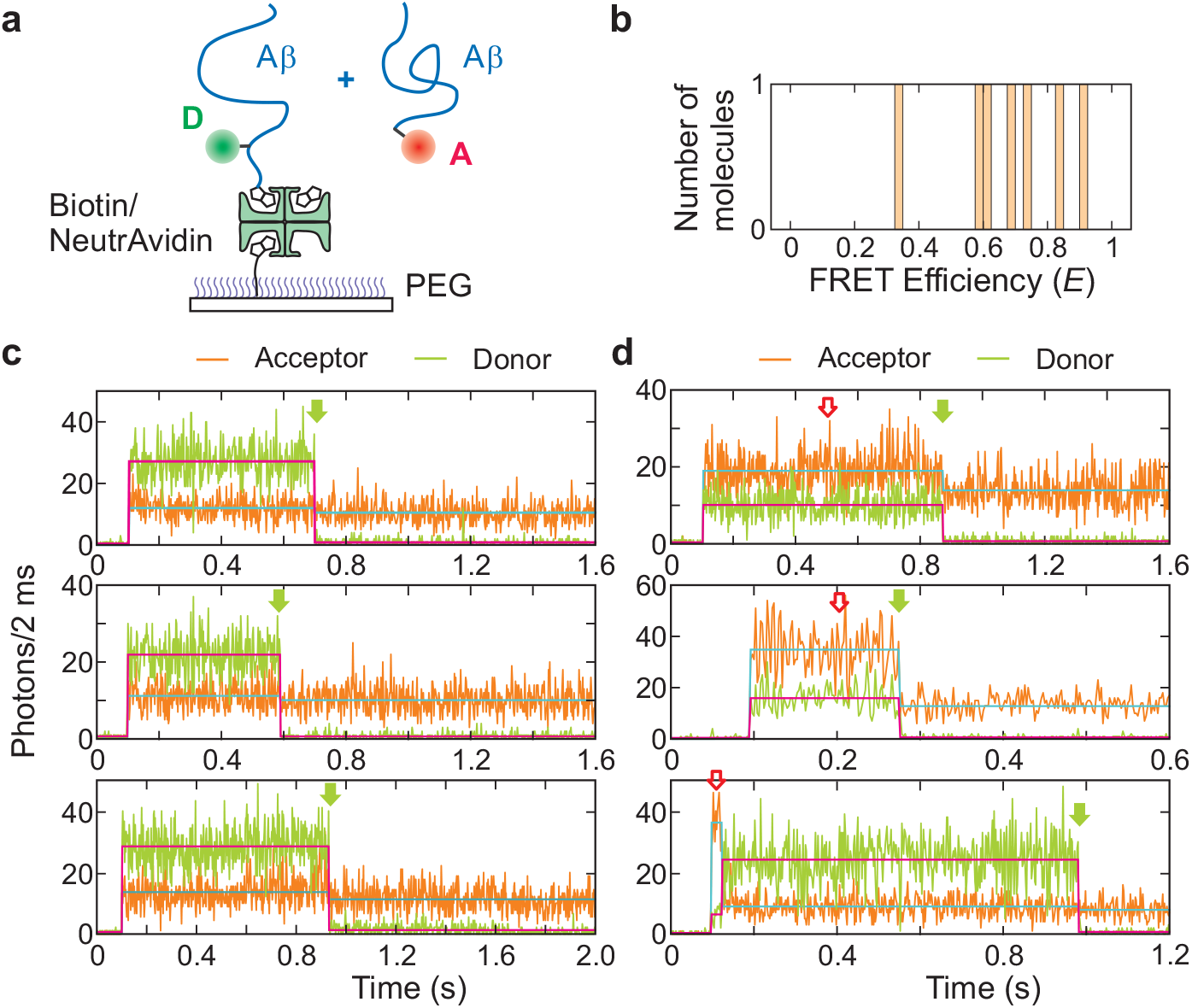
Dimerization of Aβ42. **a**, Donor (D)-labeled Aβ42 (Avi-D-Aβ42) is immobilized on a polyethylene glycol-coated glass surface and incubated with acceptor (A)-labeled Aβ (A-Aβ42) at 300 nM. **b**, FRET efficiency histogram of the dimer. FRET efficiency values were corrected for background, donor leak, and *γ*-factor^11–13^. **c**, Representative donor and acceptor fluorescence trajectories (2 ms bin time) of donor-labeled Aβ42 monomers. Green arrows indicate photobleaching of the donor. The background acceptor signal after donor bleaching results from acceptor fluorescence of A-Aβ42 in solution that is excited by the donor excitation laser. Magenta and cyan horizontal lines indicate the mean photon count rates of donor and acceptor segments, respectively. **d**, Representative donor and acceptor fluorescence trajectories (2 ms bin time) of Aβ42 dimers. Open red arrows indicate the dimer segments.

The single-molecule trajectories shown in Fig. 1c are those of monomers because when the donor fluorescence signal disappears (i.e., photobleaching, green arrow), there is only a very small change in the acceptor signal. The constant, high-level background acceptor signal results from excitation of acceptors attached to Aβ in solution (300 nM) by the donor excitation laser. The very slightly higher acceptor intensity before donor bleaching results from leakage of donor photons into the acceptor channel (~ 6%). In contrast, the trajectories in Fig. 1d show large changes in the acceptor signal upon donor bleaching (green arrow). This change in acceptor emission indicates there is energy transfer from a donor of an immobilized Aβ (Avi-D-Aβ42) molecule to an acceptor attached to a monomer that was free in solution, but has bound to the immobilized Aβ to form a dimer. The possibility that oligomers consisting of more than one acceptor-labeled Aβ (A-Aβ42) are detected is low because the concentration of larger oligomers at low Aβ concentration and prior to aggregation is expected to be much lower than the dimer concentration. We detected only 7 trajectories that begin with the dimer state among 2,758 immobilized molecules (All the dimer trajectories observed in this experiment are plotted in Supplementary Fig. 2). From the fraction of the dimer and the incubation concentration of 300 nM, we obtain the dimer dissociation constant of 240 (± 90) μM (labeling efficiency is assumed to be 100% based on the UV-vis absorption measurement). Note that the dissociation constant of an indistinguishable dimer (e.g. unlabeled Aβ42 dimer) is twice the dissociation constant of a distinguishable dimer (e.g., dimer of Avi-D-Aβ42 and A-Aβ42)^14^. The FRET efficiencies of dimers vary widely ranging from 0.3 to 0.9 (Fig. 1b), indicating that dimer structures are heterogeneous.

This weak interaction between Aβ monomers might possibly result from the bulky fluorophores attached to the N-terminus of Aβ. However, we verified that the aggregation features of dye-labeled Aβ42 are not so different from those of unlabeled Aβ42 (Supplementary Figs. 3 and 4). First, the aggregation time is similar between labeled and unlabeled Aβ42 (Supplementary Fig. 3a and b). In addition, 5% addition of the sonicated sample after aggregation of labeled Aβ42 eliminates the lag phase, indicating that dye-labeled Aβ42 can be an aggregation seed similar to unlabeled Aβ42 (Supplementary Fig. 3c). Furthermore, electron microscope images of fibrils of Alexa 594-labeled Aβ42 are indistinguishable from those of unlabeled Aβ42 (Supplementary Fig. 4). Together, these results strongly suggest that fluorophores interfere minimally with oligomerization and aggregation.

### Formation of stable oligomers

Step-by-step characterization of aggregation beyond the dimer is difficult because of the weak interaction between monomers. Instead, we performed an experiment that directly detects and characterizes stable, soluble oligomers appearing in the middle of the aggregation process, some of which can be potential seeds for fibril formation. Soluble oligomers have been used in various biological toxicity experiments^7,8^. In our experiment, donor-labeled Aβ42 was mixed with a large excess of acceptor-labeled Aβ42 (1:10 or 1:100 ratio) and incubated in a plate reader at 37°C (Fig. 2a). At several time points during the incubation, a small amount of sample was taken and diluted to 50 pM - 1 nM for single-molecule free-diffusion experiments. In this experiment, molecules are not immobilized, but freely diffuse and emit a burst of fluorescence photons when they briefly reside in the focal volume. Due to the very low concentration, it is possible to selectively detect single oligomeric species that survive during the measurement time of ~ 1 h. This combination of plate reader monitoring and free-diffusion experiment is crucial for sampling oligomers at particular time points during the course of aggregation given the large experiment-to-experiment variation in the aggregation time, which would be very difficult to control. For example, the aggregation time of blue trace is almost twice that of the other two in Fig. 2b (1:10 ratio, left). If sampling is done at the same time point, e.g., 1.2 h, it is before the aggregation of the blue trace and during the aggregation for the other two experiments.

**Fig. 2.**
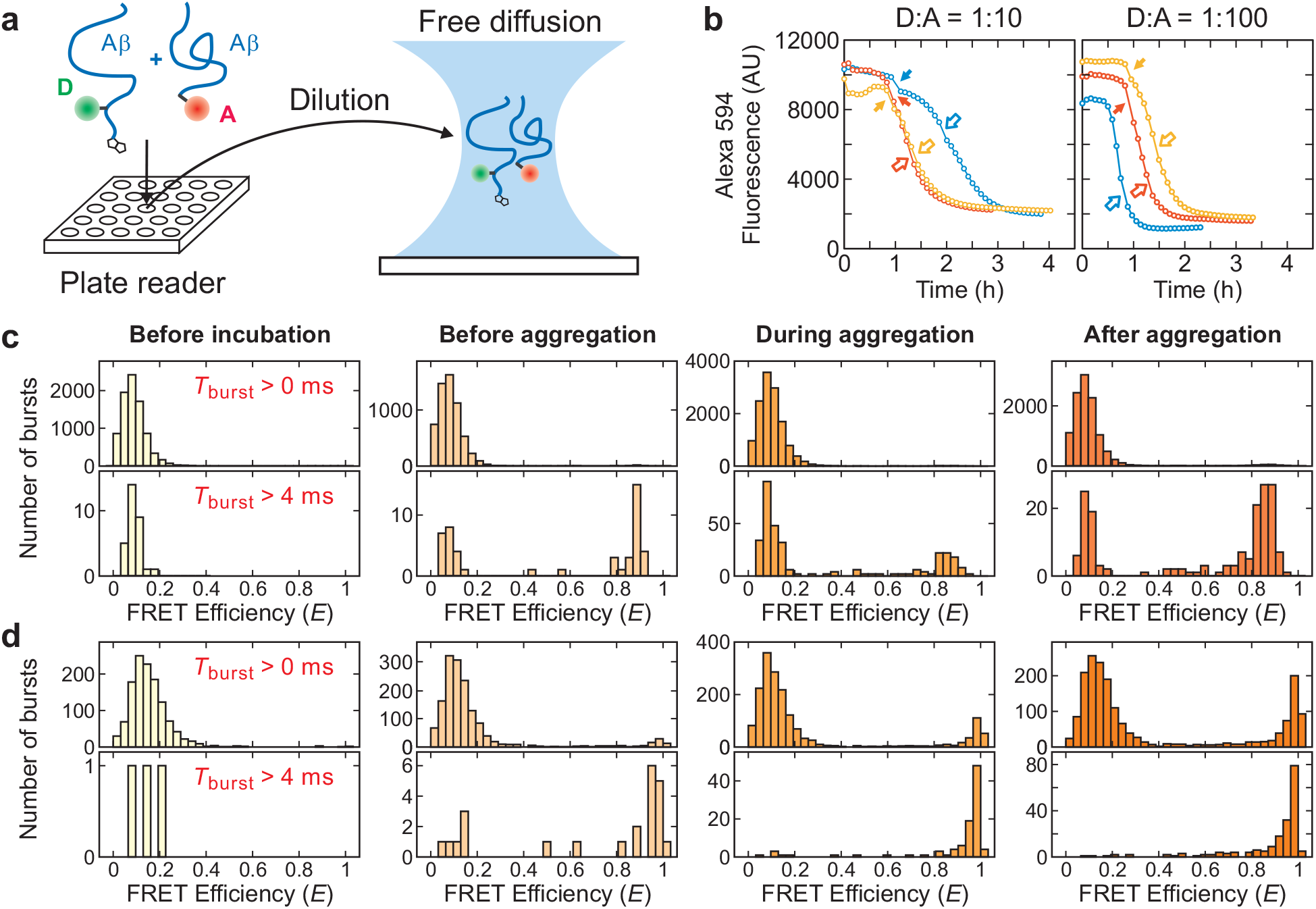
Detection of stable Aβ42 oligomers. **a**, Avi-D-Aβ42 and A-Aβ42 are mixed in 1:10 or 1:100 ratios (the concentration of A-Aβ42 is 1 μM) and incubated in a plate reader at 37°C. 0.5 - 5 μL of solution was collected from the plate reader and diluted to 50 pM – 1 nM (monomer of Avi-D-Aβ42) for the free diffusion experiment at 4 time points: before incubation, before aggregation, during aggregation, and after aggregation determined from the fluorescence intensity profile in **b**. **b**, Acceptor (Alexa 594) fluorescence intensity profile during aggregation at three different experiments (different colors). As the aggregation proceeds, the fluorescence signal decreases because the concentration of the monomer in the detection volume decreases due to the sedimentation of fibrils. The time points when the sample was collected are indicated by closed (before aggregation) and open (during aggregation) arrows. **c, d**, FRET efficiency histograms of fluorescence bursts with the average count rate greater than 40 ms-1 and with two different criteria for burst duration (Upper: *T*_burst_ > 0 ms; Lower: *T*_burst_ > 4 ms) for the experiment with a mixing ratio, **c**, D:A = 1:10 (blue trace in **b**) and **d**, D:A = 1:100 (red trace in **b**).

We performed this experiment at four different time points: before incubation, before aggregation (~ 1 h after incubation, filled arrows in Fig. 2b), during aggregation (open arrows in Fig. 2b), and after aggregation. In the experiment before incubation with a 1:10 mixing ratio, the fluorescence bursts are almost entirely donor-only, indicating that only monomers are present (Fig. 2c). Before aggregation starts, several oligomer bursts appear at high FRET efficiency of 0.85. Fibril formation might occur from some of these oligomers. The relative population of this high FRET oligomeric species increases as the aggregation proceeds. However, the overall population of oligomers is very low compared to the monomer even during and after aggregation, suggesting that oligomers have mostly dissociated into monomers upon dilution to 50 pM – 1 nM or that stable oligomers are rare.

To obtain structural information from the FRET efficiency measurements, we first wanted to determine the size of the detected oligomers. Fig. 2c shows the FRET efficiency histograms at two different burst duration cutoffs for the same data. The upper panels include all the bursts containing 30 photons and more. The bottom histograms display the bursts longer than 4 ms. Typically, the residence time of small proteins in the focal volume is several hundred μs. Therefore, the number of monomer bursts decreases significantly with the increasing duration cutoff (see Supplementary Figs. 5 and 6 for 2 ms cutoff). On the other hand, relatively more oligomer bursts survive at 4 ms cutoff. This indicates that diffusion of the detected oligomers is much slower than that of the monomer due to the larger size. Supplementary Fig. 5 shows two-dimensional plots of the photon count rate vs. burst duration of individual bursts. The shape of the distribution for the monomer and oligomers is clearly different. In addition, there are many more bursts with high count rates and longer burst durations (> 10 ms) for oligomers compared to the monomers, also indicating much large size of oligomers. Supplementary Fig. 7a and b shows our previous study of the tetramerization domain (TD) of p53 as a comparison. This peptide consists of 42 residues and the donor-labeled protein construct has the same biotin accepting sequence (AviTag, Supplementary Fig. 1) followed by the linker sequence identical to that of Avi-C-Aβ42 in this study. The oligomerization of TD stops at the tetramer, and there is no long-duration burst from the tetramer (*E* ~ 0.8, Supplementary Fig. 7b). This comparison shows the detected Aβ42 oligomers contain many more than 4 monomers.

The experiment with a higher donor:acceptor ratio (1:100) (Fig. 2D and Supplementary Fig. 6) shows very similar trends with the 1:10 ratio experiment. The only difference is that the FRET efficiency of oligomers is higher (*E* > 0.95) in 1:100 ratio than in 1:10 ratio (*E* = 0.85). This difference indicates that the oligomer size is much larger than a decamer. Most of the oligomer bursts would contain at least one donor-labeled Aβ42 (There are a relatively small number of bursts by acceptor only oligomers due to direct acceptor excitation in the 1:100 ratio experiment. See Supplementary Fig. 8c and d). If the size is smaller than a decamer, the majority of oligomers would have only one donor, which would not be changed much by increasing the donor/acceptor ratio. In this case, the mean FRET efficiency of the oligomers would be similar in 1:10 and 1:100 ratio experiments. On the other hand, if the size of oligomers is much larger than a decamer, there would be more than one donor in oligomers from 1:10 ratio experiment and the number of donor in oligomers will decrease in the 1:100 ratio experiment, which would result in more energy transfer from the donor to nearby acceptors. Therefore, the FRET efficiency from 1:100 mixing ratio will be higher than that from 1:10 mixing ratio. Fawzi *et al*. have shown that only monomers and protofibrils are detected by NMR^15^. Recently, Barnes *et al*. have also observed the formation of only the large-size oligomers (> 80-mer) after pressure-jump using NMR at high protein concentrations^16^. Our observation of large stable oligomers is consistent with these previous studies.

### Aggregation

The information that can be obtained from the experiments on dimerization and stable oligomer formation is limited primarily because of the weak monomer interactions and the low population of oligomers, which is unexpected given the previous studies of isolation of various forms of oligomers and their usage in biological experiments^7,8^. Since the size of stable oligomers is large, we designed an experiment to detect large oligomers along with aggregation (fibril elongation) using fluorescence lifetime imaging (FLIM). In this experiment (Fig. 3a), donor-labeled Aβ42 is immobilized and incubated with acceptor-labeled Aβ42 similar to the dimerization experiment, but at a slightly higher concentration of A-Aβ42 (500 nM) and for much longer incubation times. Immediately after the start of incubation at room temperature, 16 (4 × 4) – 36 (6 × 6) 10 × 10 μm^2^ regions of the sample were sequentially scanned. After completion of one set of scans, the stage was moved back to the first region and scans were repeated. Since scanning 36 regions takes about 50 min, for example, 24 repetitions produce 36, 20 hour-long movies (~ 50 min frame rate) of fluorescence intensity and lifetime of oligomers and fibril elongation (see Supplementary Videos 1 – 3). Fig. 3b and c show snapshots of Supplementary Videos 1 and 2 that capture various features of large oligomer formation and fibril elongation. First, oligomer formation or fibril growth from immobilized donor-labeled Aβ42 was not observed. This finding is expected because the number of immobilized molecules is so much smaller than the number in solution, so the chance of stable oligomer formation originating from immobilized molecules is extremely low. (Also, recall from Fig. 1 that even dimerization is very rare) Consequently, all detected oligomers, as well as short and long fibrils, are those that sedimented from the solution. These oligomers and fibrils containing only acceptor fluorophores (Alexa 594), which emit fluorescence after direct excitation (no energy transfer from the donor) at the donor excitation wavelength, 485 nm. The appearance of fibrils on the immobilization surface does not result from permanent sticking because some fibrils suddenly disappear (dissociation from the surface) and one end of the fibril moves for some fibrils over time.

**Fig. 3.**
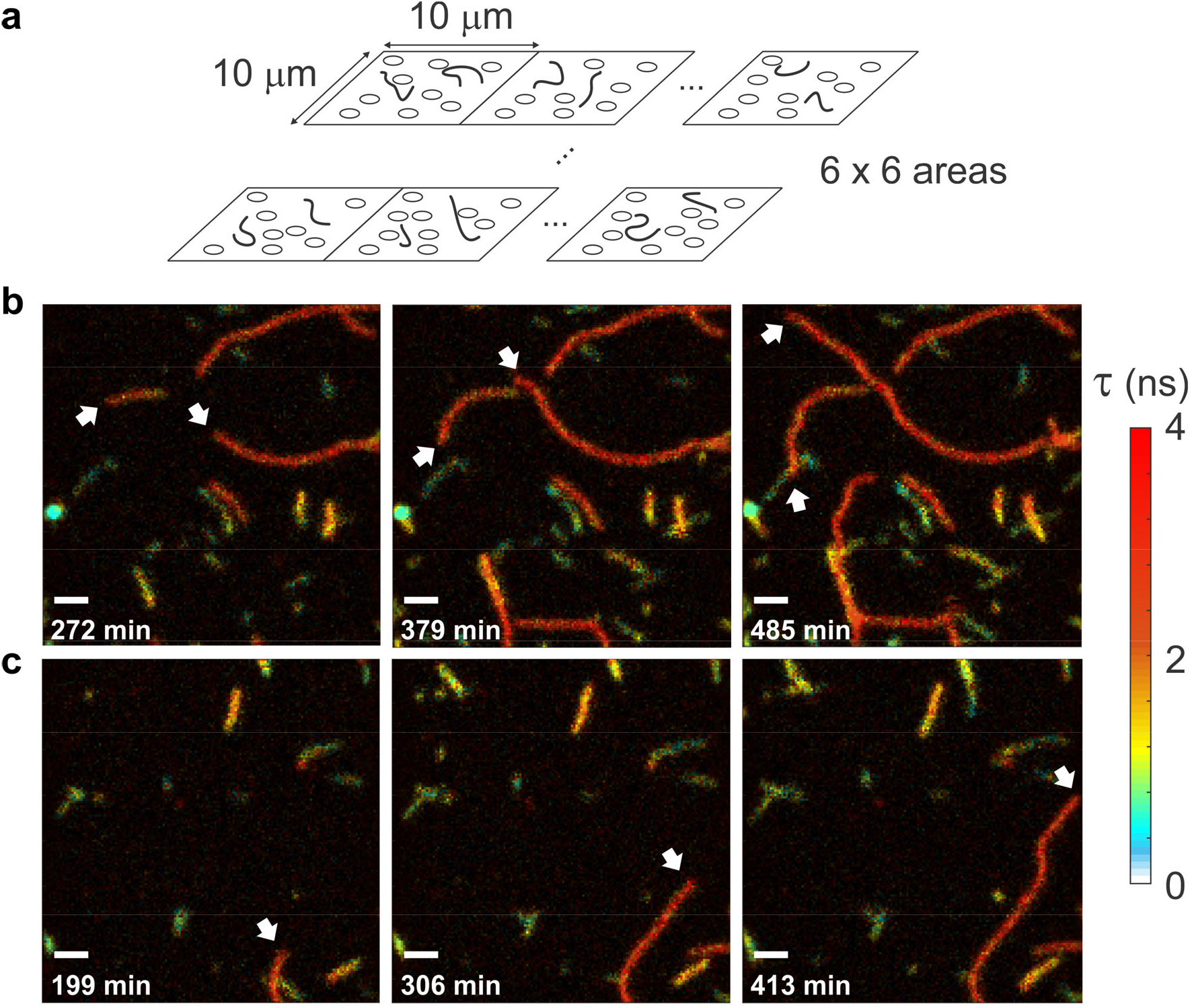
Monitoring of Aβ42 aggregation by fluorescence lifetime imaging. **a**, A PEG coated surface was incubated with 500 nM of A-Aβ42 (see Fig. 1a). Immediately after starting incubation at room temperature, 16 (4 × 4) – 36 (6 × 6) 10 × 10 μm^2^ regions are sequentially scanned. After completing one set of scans, the stage is moved back to the first region and the scans are repeated. **b, c**, Three snapshots of Supplementary Videos 1 (**b**) and 2 (**c**). Fast growing ends are indicated by white arrows. Incubation times are indicated with a 1 μm scale bar on the lower left corner of each image. Fluorescence lifetime (τ) images were masked by count rates and smoothed using Total variation denoising^17^.

The videos and snapshots in Fig. 3 show that fibril elongation is highly heterogeneous. There are relatively short fibrils, which do not grow or grow slowly (green/blue). These molecules could be large nonfibrillar oligomers or protofibrils^18,19^. In addition, there are long fibrils that elongate much faster (orange and red). Overall, the fluorescence lifetime is shorter than that of the acceptor of the monomer (3.67 ns), which can be explained by self-quenching of fluorescence when dyes are placed too close to each other^20^. Interestingly, the fluorescence lifetimes of short, slowly-growing fibrils (green and blue) are shorter than those of long, fast-growing fibrils (orange and red). Different lifetimes indicate that the structure (i.e., arrangement of monomers) is different for short, slowly growing and long, fast growing fibrils. In addition, for the fast-growing fibrils, elongation is often much faster from one of the two ends. This polarized fibril growth has been observed experimentally^21–23^ and in molecular dynamics simulations^24^.

### Individual fibril analysis

More detailed and quantitative aggregation features can be obtained from characterization of individual fibrils. For this analysis, it is required to identify and separate individual fibrils and follow their changes over time. However, as the aggregation proceeds, existing fibrils grow and new fibrils appear due to sedimentation, and they start to overlap (Fig. 4a). In many cases, it is unclear how to split the fluorescence intensity of the overlapping region into different fibrils in a single image. To solve this problem, we developed and used a deep neural network that is trained to distinguish the growth of existing individual fibrils and appearance of new fibrils by tracking the history of the entire FLIM movies. Starting from the easiest problem, the initial frames where fibrils (i.e., oligomers) don’t overlap, the deep neural network builds up information to track and characterize overlapping fibrils in later frames iteratively (see Methods and Supplementary Figs. 9 and 10). Using the deep neural network, we analyzed 3,893 image frames from 5 experiments and characterized 179,176 images of 15,004 individual fibrils.

**Fig. 4.**
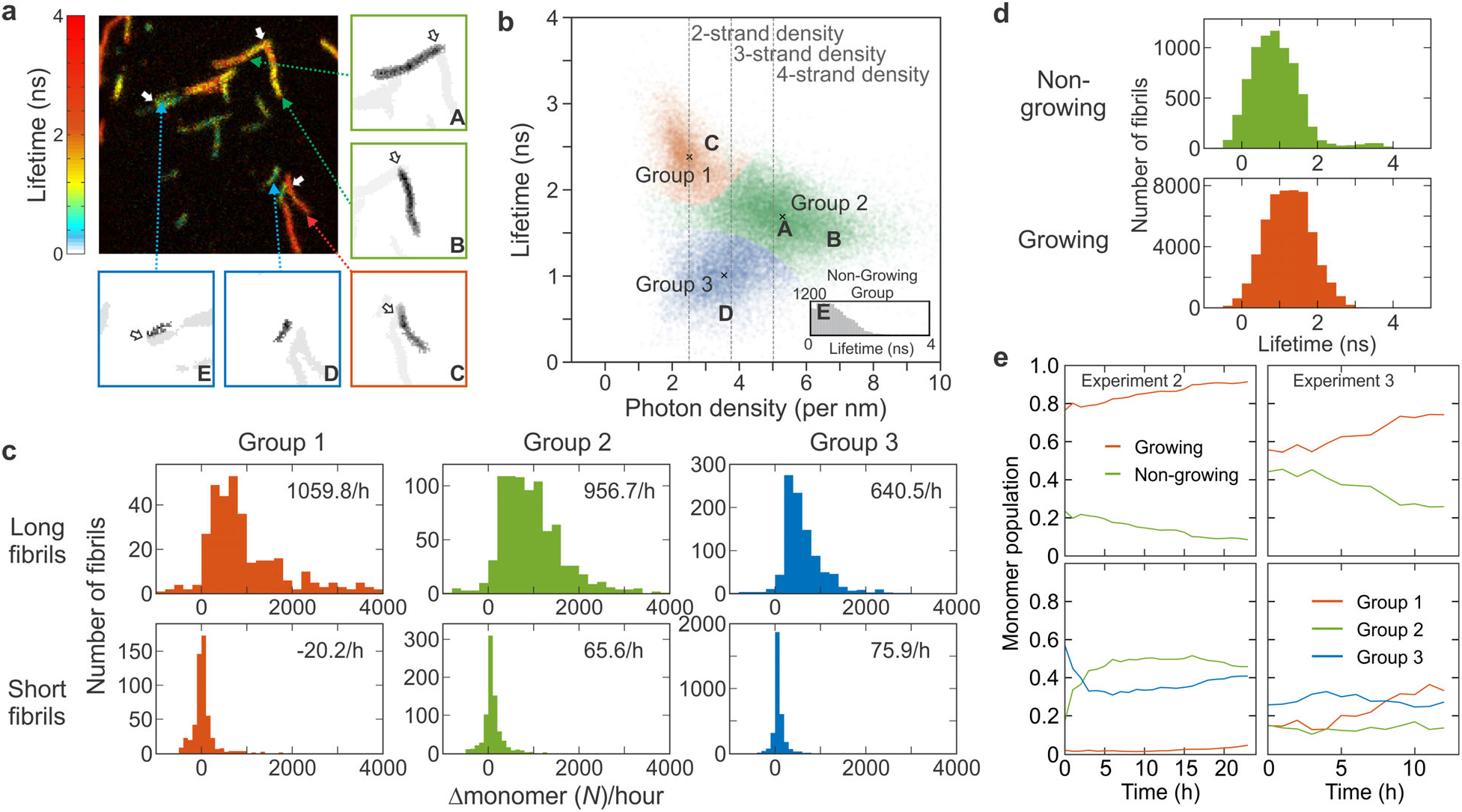
Individual fibril analysis. **a**, Individual fibrils in an image at time *t* are separated using deep learning (see Methods and Supplementary Figs. 9 and 10). Five individual fibrils with different characteristics (A – E) are shown in the sub-regions on the right and bottom side of the image as examples. Each sub-region consists of only one fibril (black), but nearby fibrils (grey) are also shown for comparison. Red, green, and blue squares of the sub-region indicate fibril group 1, 2, and 3 in **b**, respectively. White arrows indicate the overlapping regions of fibrils before separation. **b**, Fluorescence lifetime of Alexa 594 attached to Aβ42 vs. photon density (number of photons per unit length, nm). The distribution of individual fibrils with a measurable length (i.e., twice larger than the PSF size) are clustered into three groups (three different colors). An x mark indicates the center of each cluster. Fibrils belonging to the non-growing group (see text for the definition), which are often smaller than the PSF size, are characterized only by the fluorescence lifetime (inset). Letters A – E indicate the locations of the five fibrils on the plot. **c**, Growth rate of long and short fibrils of the three fibril groups. **d**, Fluorescence lifetime distributions of non-growing and growing fibrils. **e**, Time-dependent changes of the fractions of monomers in non-growing and growing fibrils (upper) and changes of three fibril groups of growing fibrils (lower) from experiment 2 and 3. See Supplementary Fig. 14 for the results from other experiments.

Fig. 4 illustrates separation of fibrils and quantitative characterization of the heterogeneity of aggregation. Five fibrils A – E are shown as examples. After separation, the lengths of these fibrils are measured and the number of photons and fluorescence lifetimes are calculated from the photons and their delay times (i.e., photon arrival times after pulsed laser excitation) collected in those fibril regions, which results in two-dimensional (2D) plot of fluorescence lifetime vs. photon density (i.e., number of photons per unit length), which is proportional to the monomer density. As seen in Supplementary Videos and in Fig. 3, there are large variations in the fluorescence lifetime and intensity. By clustering this 2D data, we classified fibrils longer than 500 nm, which is about twice the size of the point spread function (PSF, 2σ = 247 nm), into three groups using Gaussian mixture models^25^ (see Methods). Based on their lifetimes, fibrils in group 1, 2, and 3 appear in red (fibril C), yellow (fibril A, B), and green/blue (fibril D, E), respectively, in the image (Fig. 4a). Using photon density and fluorescence lifetime, it is also possible to estimate the number of strands in a fibril (i.e., polymorphism) from the length of a fibril and the number of monomers in it (see Methods for the calculation). Fig. 4b shows that group 1, 2, and 3 fibrils consist of 2, 4, and 3 strands, respectively. In addition, there are non-growing fibrils (fibril E in Fig. 4a), the majority of which are shorter than the measurable size (i.e., twice the PSF size). Since the density cannot be defined, these fibrils (or oligomers) are characterized and classified into three groups based on their lifetime distances from the average lifetimes of the fibril groups (group 1: τ > 2.04 ns, group 2: 1.35 ns < τ ≤ 2.04 ns, group 3: τ ≤ 1.35 ns) (inset in Fig. 4b).

Figure 4c - e show detailed statistics of fibril elongation analysis that reveals highly heterogenous aggregation features. We first compared the average growth rate (increase of the number of monomers per hour) of the long (i.e., measurable) and short (i.e., not measurable) fibrils of different groups (Fig. 4c). The growth rate of the short fibrils is very slow for all three groups, indicating these fibrils are classified as mostly non-growing fibrils. In addition, the majority of short fibrils belong to group 3 with short fluorescence lifetimes (compare the height of the histograms in Fig. 4c), which is consistent with the observation from example movies (Fig. 3). Among long fibrils, the growth rate of group 3 is also the lowest. The growth rate of group 1 and 2 are similar, but the number of strands of group 2 is twice as many as that of group 1 (Fig. 4b). Therefore, the apparent growing speed of group 1 in terms of length in movies looks twice as fast as that of group 2. In addition, there is a long tail in the group 1 distribution (Fig. 4c), indicating the presence of extremely fast-growing fibrils (colored in red) as seen in Fig. 3. The variation of the growth rate distribution of growing fibrils over different experiments is not large for all three groups (Supplementary Fig. 11). The maximum size of fibrils that is reached at the end of the experiments ranges widely between 1,000 and 50,000 monomers (Supplementary Fig. 12).

We define non-growing fibrils as the fibrils with an average growth rate slower than 5^th^ percentile of the growth rate distribution of the long fibrils, which corresponds to 128 monomers/h (63.4 photons/h). Fig. 4d compares the characteristics of growing and non-growing fibrils. First, the average lifetime of non-growing fibrils is shorter than that of the growing fibrils, consistent with the visual characterization of the movies (Fig. 3). The distribution varies depending on the sample batches. Supplementary Fig. 13 shows the comparison of the lifetimes of non-growing and growing fibrils from 5 experiments with two different sample batches (two different expressions of Aβ42 and labeling). The data clearly shows that there are more group 3 fibrils in both non-growing and growing fibrils with short fluorescence lifetimes in batch 2.

In addition to this overall distribution and classification, it is important to characterize how these different fibril groups change over time as aggregation proceeds. Fig. 4e compares the evolution of the population of non-growing and growing fibrils of two of five experiments (experiment 2 and 3 using batch 1 and 2, respectively). Supplementary Fig. 14 shows all five experiments. In general, the fraction of growing fibrils increases because of their faster growth rates. However, at the beginning of aggregation, the fraction of the growing fibrils varies widely between 40 – 80%. The changes of the fraction of different fibril groups are more diverse. Overall, group 3 is dominant in the non-growing fibril group (Supplementary Fig. 14b). On the other hand, for growing fibrils, group 3 dominates at the beginning of aggregation, but other groups catch up at later times (Supplementary Fig. 14c). In batch 1, the fraction of group 1 fibrils is very low over the entire time course, whereas both group 1 and 2 increases with time in batch 2 data.

### Heterogeneous secondary nucleation from oligomers

Since it is possible to follow the growth of each individual fibrils, we back-tracked the growth of fibrils with a measurable length to identify the fibril group at their first appearance (i.e., origin) as an oligomer (or protofibrils, < 500 nm). Fig. 5a shows this distribution. Group 1 and 3 fibrils originate mostly from oligomers of their own groups. However, the majority of group 2 fibrils come from group 3 oligomers when they appear at the beginning of aggregation (0 – 1 h), which is replaced by the same group 2 when the oligomers appear at later times (Fig. 5a, middle row). This apparent interconversion between different groups may support the mechanism of the aggregation seed formation by conformational conversion^26,27^. However, abrupt structural conversion of large oligomers (larger than 100-mer) would be highly improbable because many monomers need to almost simultaneously convert conformations into the same structure. Instead, we interpret this apparent interconversion of groups as the formation of a new fibril (nucleation) on the surface of an oligomer with a different structure. In this case, as a fibril grows, the group identity will change gradually from one group (original oligomer) to another (new fibril) as new monomers with a different structure group are added over time. Indeed, Fig. 5b shows gradual changes of the fluorescence lifetime distribution, supporting this mechanism. The lifetime distributions of the frame immediately before (blue) and after (orange) the identified transition (top, +/−1) from group 3 to group 2 in lifetime trajectories are close and overlap, which are located in the middle of the average lifetimes of group 2 and group 3 fibrils (vertical dashed lines). However, the distributions 5 – 6 frames away from the transition interval are separated more and become closer to the average values. This nucleation is similar to the secondary nucleation mechanism^28,29^, but different because the nucleation occurs on the surface of oligomer/protofibril rather than long fibrils. In addition, the structure of a newly-formed fibril is different from the structure of the original oligomer/protofibril. Therefore, we call this heterogeneous secondary nucleation. (see Fig. 6).

**Fig. 5.**
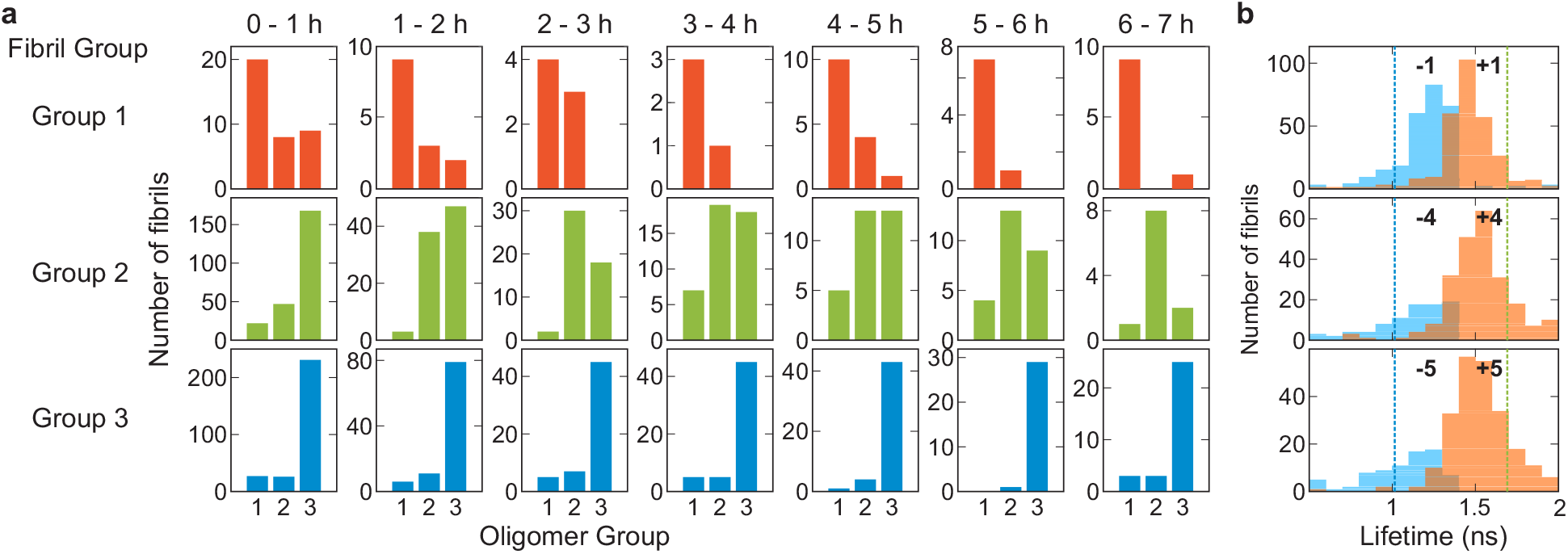
Heterogeneous secondary nucleation from oligomers. **a**, Oligomer origin of the three fibril groups (three rows). The growth of individual fibrils of the three groups was back-tracked to identify the group of oligomers or short fibrils (< 500 nm) at their appearance during the time period of 0 – 7 hours. The majority of group 1 and 3 fibrils originate from the oligomers of their own groups, whereas the majority of group 2 fibrils originate from group 3 oligomers when they appear at the beginning of aggregation (0 - 1 h), which is replaced by group 2 oligomers at later times. **b**, The distribution of fluorescence lifetimes of fibrils before (blue) and after (orange) the transition from group 3 to group 2 at three different frame separation from the transition interval: (upper) ± 1 frame, (middle) ± 5 frames, and (bottom) ± 6 frames. Vertical dashed lines show the average fluorescence lifetimes of group 2 (green) and group 3 (blue).

**Fig. 6.**
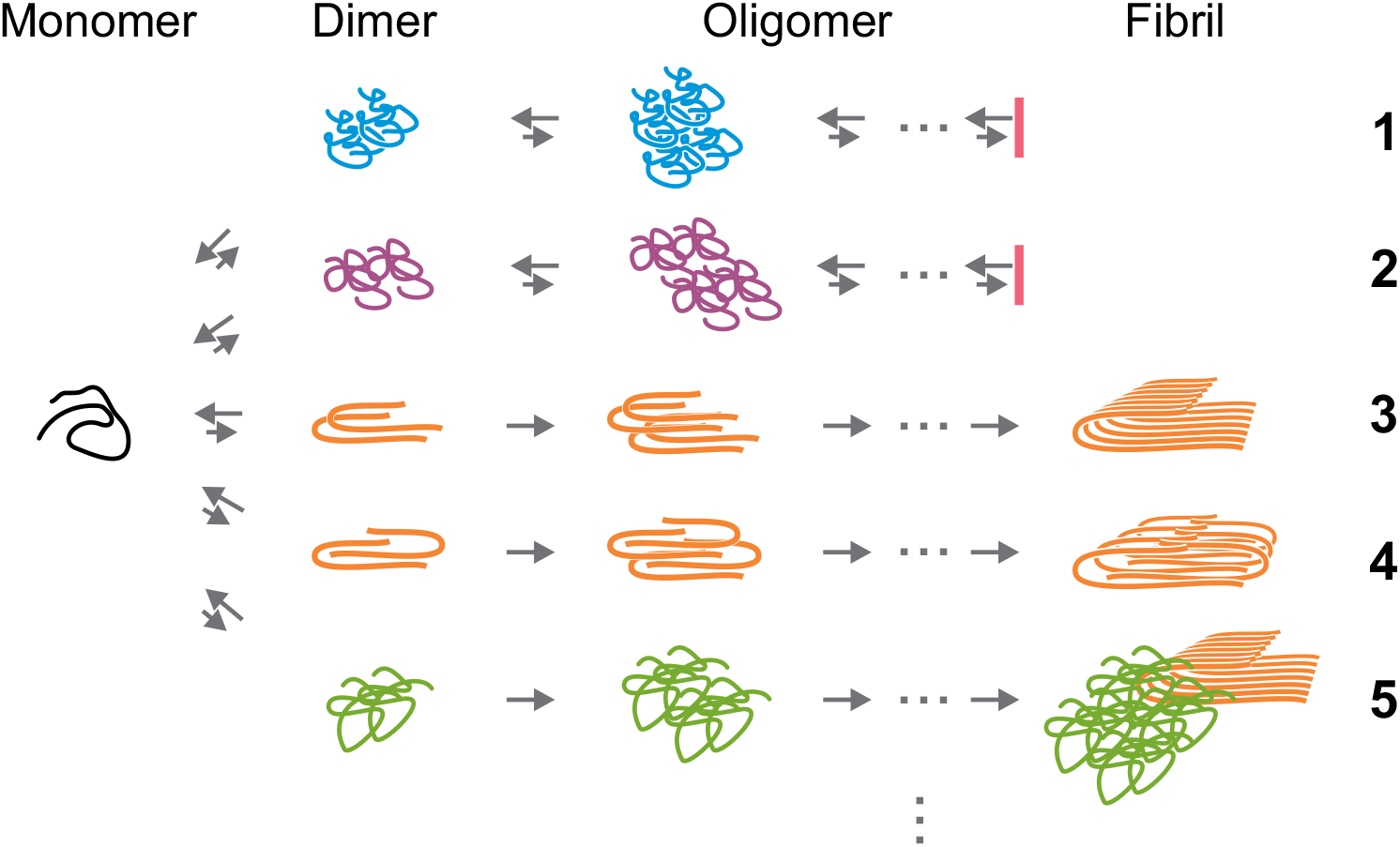
Model of heterogeneous Aβ42 oligomerization and aggregation. Initially, monomers form dimers with different structures, which are followed by diverse oligomerization and aggregation pathways. There are pathways (1 and 2) in which the assembly stops in oligomeric stages with the possibility of backtracking^34,35^ and pathways (3 and 4) that lead to fibril elongation. There are fibrils growing on the surface of oligomers with different structures (heterogeneous secondary nucleation, pathway 5).

## Discussion

The major problem in characterizing Aβ42 aggregation is that the experimental results vary widely depending on experimental methods and are often irreproducible. The size of stable oligomers, for example, which have been reported to show biological toxicity, ranges from dimers to large protofibrils^3–6,30–32^. However, there is almost no information on the conformation, stability, and relative population of different oligomers. As for the aggregation mechanism, the most quantitative model that explains the long lag time and its non-linear concentration dependence is the secondary nucleation model^28,29^ of Ferrone *et al*.^33^ that accounted for the lag phase and large nucleus size for the aggregation of sickle hemoglobin.

Our experimental results directly demonstrate the heterogeneity in the oligomerization and aggregation, which varies widely over samples prepared at different times (different batches) as well as experiments performed using the same batch of sample. These results suggest that a wide variation in experimental results using different protocols and techniques is fully expected due to the heterogeneous nature of Aβ42 aggregation. In one experiment, one set of oligomerization and aggregation pathways are preferred, while under even slightly different conditions, a different set of pathways is preferred. Our first finding to support this conclusion is that dimerization is mediated by weak monomer-monomer interactions (*K*_d_ = 240 μM), which results in diverse dimer conformations as evidenced by differences in FRET efficiency (Fig. 1b). We then used a combination of plate reader and single-molecule free-diffusion experiments to detect soluble, stable oligomers, some of which could be aggregation seeds and potentially toxic species as demonstrated by various studies^7,8^. The concentration of these oligomers is very low. Finally, we showed that aggregation is highly diverse in terms of the length, elongation speed, and structure of fibrils (Figs. 4 and 5 and Supplementary Figs. 11 – 14).

A model of heterogeneous oligomerization and aggregation based on these observations is shown in Fig. 6. Initially, monomers form dimers with various conformations. These dimers can grow into oligomers of larger size, but many of these oligomers may stop growing into long fibrils (i.e., nonfibrillar oligomers, pathway 1 and 2 in Fig. 6 as examples). These pathways correspond to the formation of short and slowly- or non-growing fibrils in Fig. 4 such as group 3 oligomers. Some of the oligomers would grow into long fibrils, which correspond to fast-elongating fibrils in the experiment (pathway 3 and 4 in Fig. 6). Oligomers that do not grow into fibrils may very well be the oligomers discovered in simulations by Wolynes and coworkers^34,35^ that require dissociation into monomers to proceed to fibrils, a process called “kinetic backtracking.”

It has been proposed that Aβ forms oligomers without β-structures first, which is followed by the structural conversion to cross-β structures that promote the fibril elongation^26,27^. If this happens, it may appear as transitions between different groups as shown in Fig. 5. However, abrupt structural conversion of large oligomers (larger than 100-mer) would be highly improbable because many monomers need to almost simultaneously convert conformations. We therefore propose another mechanism in which fibril elongation occurs on the surface of (potentially) non-growing oligomers (group 3). The gradual changes in fluorescence lifetime support this mechanism (Fig. 5b). This is similar to the secondary nucleation mechanism, in which the nucleation occurs on the surface of existing fibrils, but in our case, the structure of the fibril is different from that of the parent oligomer (or short fibril). To distinguish our observation from the original mechanism, we call this process “heterogeneous secondary nucleation.” (pathway 5 in Fig. 6) In fact, we have not directly detected the change corresponding to the original secondary nucleation mechanism that may appear as growth of a very short fibril from the middle of another fibril. However, this does not reject the original mechanism because small nuclei would not be detected in our experiment and oligomers that nucleate on parent fibrils may rapidly detach^36^. In addition, our observation does not exclude the possibility of conformational conversions between small oligomers, which cannot be detected in our experiment due to the low photon count rates from small oligomers.

The observation of highly heterogeneous pathways that may be sensitive to the environment suggests that oligomerization and aggregation pathways *in vivo* may be quite different from those observed *in vitro*. The physiological concentration of Aβ42 is much lower than 1 μM, and, therefore, oligomerization and aggregation by Aβ42 alone would be very improbable. Since Aβ can interact with many other cellular components non-specifically, these interactions may promote oligomerization, nucleation, and eventually fibril formation similar to the heterogeneous secondary nucleation that we observed. Different structures of fibrils grown from brain-derived aggregation seed supports this hypothesis^37,38^. In this case, characterization of the oligomer heterogeneity in the context of cellular toxicity will be critical to understand the disease mechanism and to discover targets for drug therapy.

## Methods

Details of the expression, purification, and dye labeling of Aβ42, data collection using single-molecule fluorescence experiments and fluorescence lifetime imaging, and data analysis including the development of the deep neural network are described in the Supplementary information (codes are available at https://github.com/hoisunglab/FNet).

## Acknowledgements

We thank W. A. Eaton, A. Szabo, R. Tycko, I. V. Gopich, J.-Y. Kim for numerous helpful discussions and comments and J. M. Louis for advice and suggestions on protein expression and purification. This work utilized the computational resources of the NIH HPC Biowulf cluster. (http://hpc.nih.gov). This work was supported by the Intramural Research Program of the National Institute of Diabetes and Digestive and Kidney Diseases, NIH.

## Author Contributions

H.S.C. directed research. F. M. prepared protein samples and performed single-molecule and FLIM experiments. J. Y. developed deep learning-based fibril analysis method. F. M., J. Y., and H.S.C. analyzed the data and wrote the manuscript.

## Data Availability

All data is available in the manuscript or Supplementary Figures. Additional analysis data and materials used in this study are available upon request.

## Code Availability

Custom codes for the deep neural network are available at https://github.com/hoisunglab/FNet.

## Supplementary Information

### Methods

#### Protein expression

The amino acid and DNA sequences of Aβ42 are shown in Supplementary Fig. 1. For labeling of the donor (Alexa 488) and acceptor (Alexa 594) dyes, a cysteine residue was attached to the N-terminal of Aβ42 (Avi-C-Aβ42 and C-Aβ42). To immobilize proteins on a biotin-embedded glass coverslip, a biotin accepting sequence (AviTag, Avidity LLC, Aurora, Colorado) and a flexible linker sequence were attached to the N-terminus of Aβ (Avi-C-Aβ42). All plasmids were constructed by DNA2.0 (DNA2.0, Neward, CA). To ensure the expression of biotinylated proteins, we co-expressed the BirA gene to generate sufficient biotin ligase (Avidity LLC).

We co-transformed *E. coli* strain BL-21 (DE3) (Stratagene, La Jolla, CO) with kanamycin-resistant pJ411-BirA, and carbenicillin-resistant pJ414-Aβ, for the expression of Avi-C-Aβ42. For C-Aβ42 and C-Aβ42, we transformed the bacteria with pJ414-Aβ. The expression level of the full-length protein was optimized by varying the ratio of the plasmids. The optimized condition was 0.2 μL of pJ411-BirA (50 ng/μL) and 0.2 μL of a protein construct (20 ng/μL). Co-transformed bacteria were spread on LB-agar plates with corresponding antibiotics. After incubation at 37°C overnight, 2 - 3 individual colonies were picked and inoculated in 5 mL LB broth with the same antibiotics combinations for 16 - 24 hours at 37°C with shaking at 250 rpm. Colonies grown up in liquid medium were diluted into the same medium of 500 - 1000 mL for further growth. After incubation for 3 - 5 hours, expression was induced at OD 0.6 (600 nm) with final concentrations of 1 mM IPTG, and 50 μM *d*-biotin. After overnight incubation at 25°C with shaking at 250 rpm, bacteria was harvested and spun down at 8000 g for 10 minutes using Sorvall LYNX 4000 centrifuge (Thermo Scientific, Waltham, MA). After removing the supernatant, pellets were either used for lysis right away or frozen at −20°C for future use.

#### Purification of proteins

Bacteria pellets from 500 mL LB culture were lysed in 20 mL of bacterial protein extraction reagent (B-Per, Thermo Fisher Scientific, Grand Island, NY) with 50 mM benzamidine hydrochloride, 100 μg/mL lysozyme (Sigma, St. Louis, MO), and 5 units of benzonase (Novogen, Madison, WI). The pellets were mixed and resuspended in the lysis buffer and incubated at room temperature for 30 minutes. The lysate was transferred to 50 mL spinning tubes and centrifuged at 30000 g for 45 minutes with Sorvall LYNX 4000. The supernatant was removed for electrophoresis and the pellet containing inclusion bodies were resuspended in 30 mL 1× PBS solution with 10 mM DTT and 1% Triton X-100 and sonicated three times for 20 seconds on ice using a sonicator at 100% power (Model Q55, Qsonica, Newtown, CT). The solution was then centrifuged at 30000 g for 30 minutes at 4°C. The supernatant was discarded and the remaining pellet was resuspended in the same PBS buffer used in the previous sonication step. One molar sodium chloride was added to remove DNA and RNA from pellets. The mixture was sonicated as in the previous step and centrifuged at 30000 g for 30 minutes at 4°C. Resuspension, sonication, and centrifugation were repeated in 1× PBS. The pellet containing inclusion bodies was dissolved in 5 mL of 50 mM Tris-HCl with 6 M guanidine hydrochloride (GdmCl) and 10 mM DTT and kept at room temperature overnight for complete extraction of Aβ proteins. The solution was then centrifuged at 30000 g at 4°C for 45 minutes to remove the insoluble pellet. The supernatant was collected for further purification. The supernatant was loaded on PhastSystem (Pharmacia, Baltimore, MD) gels. Gels were stained with Phastgel Blue R (Pharmacia, Baltimore, MD) then washed until protein bands were clearly shown. Aβ proteins with and without AviTag and linker appeared at 8 kDa and 5 kDa on gels, respectively, and these were the smallest proteins presenting in the inclusion body. The protein solutions (200 μL) were loaded onto the AKTA pure FPLC system equipped with a Superdex™75 10/300GL size exclusion column (GE Healthcare, Chicago, IL). The separation was run with 50 mM Tris-HCl, 4 M GdmCl solution at a flow rate of 0.8 mL/min. The fractions containing 5 kDa or 8 kDa proteins identified by Phastgel were collected and concentrated using Amicon Ultra centrifugal filters (EMD Millipore, Billerica, MA) and then subjected to the second round of FPLC purification.

#### Dye-labeling and purification

We labeled Avi-C-Aβ42 with Alexa Fluor 488 maleimide (Alexa 488, A10254, Thermo Fisher Scientific, Carlsbad, CA) and C-Aβ42 with Alexa Fluor 594 maleimide (Alexa 594, A10256, Thermo Fisher Scientific, Carlsbad, CA). Aβ (~ 0.2 mg) in 4 M GdmCl Tris buffer solution (pH 8) was concentrated to 100 μL in 6 M GdmCl using Amicon centrifugal filters and pH was adjusted to 7.0 by acetate buffer. 100 μL of protein solution was mixed with 0.1 mg of Alexa 488 or 594 pre-dissolved in 5 μL of DMSO. We incubated the mixture at room temperature overnight. Then the reaction was quenched by adding 4 μL of β-mercaptoethanol. The reaction mixture was fractionated on a Superdex™75 10/300GL size exclusion column equilibrated with 50 mM Tris-HCl, pH 8.0, 4 M GdmCl to remove the excess free dye. The peptide labeled with the dyes showed overlapping peaks of absorbance monitored at 280, 494 nm for Alexa 488, and 280, 594 nm for Alexa 594. The labeled protein concentration was determined by the absorbance at 494 nm or 594 nm measured by Cary 8454 UV-Vis spectrophotometer (Agilent Technologies, Santa Clara, CA). Purified samples were aliquoted into 10 μL and kept at −80°C for future experiments.

#### Plate reader experiment for preparation of stable oligomer mixture

We mixed donor-labeled Aβ (Avi-D-Aβ42) with 1 μM of acceptor-labeled Aβ (A-Aβ42) at the ratios of 1:10 and 1:100. The mixture (20 - 50 μL) was loaded into 96-well half-bottom none binding surface polystyrene plates (REF 3881, Corning, Kennebunk, ME) for plate reader recording (Spark, TECAN, Switzerland). To monitor the aggregation of unlabeled Aβ42, Aβ42 stock solution (30 μM) was diluted into 6 μM thioflavin T (ThT, Sigma, St. Louis, MO) in 1× PBS (pH 7.4) to the final concentration of 2 μM and volume of 50 μL. All wells were sealed with a piece of parafilm to prevent evaporation in the bottom reading mode. The aggregation was monitored at 37°C for several hours. The fluorescence signal was recorded every 5 min. Samples were excited (50 flashes) at 420 nm (ThT), 475 nm (Alexa 488), and 575 nm (Alexa 594) and fluorescence was detected at 480 nm (ThT), 530 nm (Alexa 488), and 627 nm (Alexa 594) with a 20 nm bandwidth. The focus along *z*-axis was set manually at 29500 μm and the automatic gain regulation feature was used.

For the single-molecule free-diffusion experiment (see below), 0.5 - 5 μL of the mixture was collected and diluted to variable concentrations of 50 pM –1 nM (in terms of the monomer concentration of the donor-labeled Aβ) depending on the aggregation stages (see Supplementary Figs. 5 and 6 for the final concentrations). The sample was collected at 4 different time points by monitoring fluorescence changes during the aggregation process: before incubation (BI, right after mixing donor- and acceptor-labeled proteins), before aggregation (BA, during the lag phase prior to the aggregation), during aggregation (DA), and after aggregation (AA). The time points of sample collection for the BA and DA measurements are indicated in Fig. 2b. At each time points, the plate reader was briefly paused and the sample was collected after being gently stirred with a pipet tip. The well was re-sealed and the measurement was resumed after waiting for 2 min for temperature equilibration. We also performed single-molecule experiments for fibril fragments generated by sonicating the sample after aggregation. 15 μL of the aggregated sample was transferred into a 0.5 mL Eppendorf tube and sonicated with a tabletop sonicator (8890, Cole-Parmer, Vernon Hills, IL) for 3 to 5 min. Due to the depletion of the monomer during the aggregation, the final concentration for the free-diffusion experiment varies (see Supplementary Figs. 5 and 6 for the final concentrations).

#### Electron microscopy experiment

The fibrils of unlabeled Aβ42 and A-Aβ42, prepared by incubating 5 μM proteins at 37°C overnight, were imaged with FEI Morgagni microscope, which was operated at 80 kV and equipped with an AMT Advantage HR CCD camera. The samples were adsorbed to glow-discharged carbon films on lacey-carbon-coated copper mesh grids for 1 min, rinsed with deionized water, stained with 3% uranyl acetate for 1 min, and dried in air for imaging.

#### Single-molecule spectroscopy

Single-molecule FRET experiments were performed using a confocal microscope system (MicroTime200, Picoquant) with a 75 μm diam. pinhole, a beamsplitter (Z488/594rpc, Chroma Technology), and an oil-immersion objective (UPLSAPO, NA 1.4, × 100, Olympus) ^1^. Alexa 488 was excited by a 485 nm diode laser (LDH-D-C-485, PicoQuant). Alexa 488 and Alexa 594 fluorescence was split into two channels using a beamsplitter (585DCXR, Chroma Technology) and focused through optical filters (ET525/50m for Alexa 488, E600LP for Alexa 594, Chroma Technology) onto photon-counting avalanche photodiodes (SPCM-AQR-16, PerkinElmer Optoelectronics).

In the free-diffusion experiment for the detection of stable oligomers, molecules are not immobilized, but freely diffuse and emit a burst of fluorescence photons when they pass through the laser focus. Samples were prepared in 1× PBS, pH 7.5. To prevent sticking of Aβ to the glass surface, 0.01% Tween-20 was added to the solution. To reduce photoblinking and photobleaching, 40 mM cysteamine and 100 mM b-mercaptoethanol were added to the solution. Samples were illuminated in the continuous-wave (CW) mode of the laser at 20 – 25 μW. Photons with inter-photon times shorter than 300 μs were combined into one burst and bursts with 30 or more photons were considered as significant bursts and analyzed.

In the dimerization experiment, donor-labeled Avi-D-Aβ42 molecules were immobilized on a biotin-embedded, polyethyleneglycol-coated glass coverslip (Bio_01, Microsurfaces Inc.) via a biotin (surface)-NeutrAvidin-biotin (protein) linkage^2^. After being cleaned with deionized water and dried with a stream of nitrogen, the surface was covered with Cover well (PC8R-0.5) and pretreated with 20 μL streptavidin solution (25 μg/mL) for 5 minutes. The solution was replaced with 15 μL of 100 pM Avi-D-Aβ42 solution and checked on the microscope to monitor the immobilization of molecules on the surface. After observing immobilization of a sufficient number of molecules (50 – 100 molecules per 10 × 10 μm^2^), the solution was replaced with 300 nM A-Aβ42 in 1× PBS, pH 7.5, including a cocktail of 100 mM β-mercaptoethanol, 10 mM Cystamine^3^, 2 mM 4-nitrobenzyl alcohol (NBA), 2 mM cyclooctatetraene (COT), and 2 mM Trolox^4,5^ to reduce photoblinking and photobleaching of dyes. Molecules were illuminated in the pulsed mode at 1 μW.

All experiments were performed at room temperature (22°C).

#### Fluorescence lifetime imaging (FLIM)

The donor-labeled Avi-D-Aβ42 was immobilized as described in the dimerization experiment above and incubated with 500 nM of A-Aβ42 including the same chemical cocktail. A region of 10 × 10 μm^2^ was raster scanned in the pulsed mode of the laser at 0.2 μW. The scan was repeated for 16 (4 × 4) – 36 regions (6 × 6). After finishing one round of scans (40 – 50 min), the stage was moved back to the first region and the scan was repeated, which results in movies of each region of 14 – 24 hours.

#### Deep neural network

For the analysis of individual fibrils, it is required to identify and separate fibrils and follow their changes over time. However, as fibrils grow, they start to overlap, and in many cases, it is unclear how to split the changes of photon counts in the overlapping region into different fibrils in a single image. A state-of-the-art method for semantic segmentation of touching and overlapping biological objects such as cells is a mixed 2D-3D deep neural segmentation network using object bounding boxes, which are located at specific reference points^6^. In the case of overlapping fibrils, defining such reference points and bounding boxes is impossible due to the simple shape of fibrils (no structure like a nucleus in a cell as a referencse point). Therefore, instead of segmenting fibrils directly from an image of a single frame, we exploited temporal information by comparing two consecutive image frames. We assume that there exists a correct fibril segmentation in a previous image frame, and using this segmentation it is possible to predict the segmentation in the next frame iteratively. The first assumption is always true, because at the beginning of an experiment only a few small oligomers are present and there is no overlap. To predict the next frame segmentation, we first tried U-Net^7^ which has a good performance in segmentation of biological objects. However, U-Net showed very poor performance in segmentation of overlapping regions of fibrils. We found that the nature of classification-based image segmentation deep neural networks, which classify a pixel into a certain class, introduces discontinuity in the classification probability (*p*) in a single fibril image (e.g., *p* = 1 for the region without an overlap and *p* = 0.5 for the region where two fibrils overlap). This is not appropriate for splitting fluorescence intensity of a pixel into multiple fibrils. Therefore, we developed and trained a new neural network for photon count estimation of highly overlapping transparent biological objects (Supplementary Figs. 9 and 10).

The new neural network consists of four sub-networks: 1) classification network, 2) growth prediction network, 3) background prediction network, and 4) comparison network (Supplementary Fig. 10). The overall information flow is as follows: first, the classification network encodes features from a new input image and decodes them into feature maps of different resolution. The growth prediction network encodes features of individual known fibrils from the previous image, and then decodes them together with the feature maps of the same resolution from the classification network which contains the information of the new image. The background prediction network has the same structure with that of the prediction network, but it takes the previous background image as an input and uses independent weights. The classification, the growth prediction and the background prediction networks generate single feature image outputs which stand for their prediction power for how many photons of each pixel result from new molecules, known molecules, and background, respectively. Then, the comparison network, which has 3 convolution layers and 3 activation layers, compares relative prediction powers from the prediction networks and generates photon count images of known fibrils, newly-appearing fibrils, and background.

##### Classification network and prediction network

The base structure of the classification network and the prediction networks is U-Net^7^ like deep encoder-decoder networks. The classification network takes a photon count image as an input (160 × 160 × 1, 10 pixels were padded to the original image of 150 × 150 × 1 pixels) and the growth prediction network takes photon count images of individual fibrils from the previous frame as inputs (126 images) (Supplementary Fig. 10a). The encoding blocks (convolution, batch-normalization, rectified linear unit (ReLU) activation, and max pooling) of the classification network initially encodes 4 features (160 × 160 × 4). The following encoder blocks reduce the image width and height by half and add 4 additional features. The decoder blocks are similar to the encoder blocks, but double the image width and height and reduce the number of features by 4. The decoder blocks also have skip connections from the encoder blocks of the same image size. The prediction network has the same structure as that of the classification network, but decoder blocks of the prediction network has connections from the decoder blocks of the classification network (feature communications, Supplementary Fig. 10a). The background prediction network has the same structure as that of the growth prediction network, but with independent weights.

##### Comparison Network

Each input from the classification network and the prediction networks generate single channel image output. The comparison network takes these images as an input and generates final result images by the operations of convolution, a batch-normalization, and an ReLU activation twice, followed by a convolution and a PReLU activation. The result is photon count prediction images of newly-appearing fibrils, background, and updated known fibrils in the new image frame. The result images were then normalized pixel-by-pixel so that the number of photons of a pixel of the summed result image is equal to that of the corresponding pixel of the original input image.

##### Training data generation

Using the experiment #1 data, we identified potential single fibril locations by clustering high intensity pixels. By visual inspection, we extracted 1483 single fibril movies. Training images were generated by randomly rotating, reflecting and placing fibrils with variations of photon counts by multiplying a random factor ranging from 0.5 to 1.5 to the original fibril movies. Background photons with Poissonian statistics were generated with a mean value of 3.

##### Adaptive supervised learning

A training image set that mimics the actual experimental data resulted in many incorrect segmentations for overlapped and fast-growing fibrils, probably due to the relatively small fraction of photons resulting from those rare events. Therefore, we employed an adaptive learning strategy to enhance learning. In this method, new training data was generated based on the previous training result by changing 4 parameters: 1) number of initial fibrils, 2) number of newly appearing fibrils, 3) length distribution of fibrils, and 4) growth speed (acceleration). For the length distribution of fibrils, we categorized fibrils into three groups, short (number of pixels < 200), mid (200 ≤ number of pixels < 400), and long (number of pixels ≥ 400) fibrils, and adjusted relative populations of them. For the growth speed acceleration, a certain number of frames were omitted when generating a next frame image. For each generation, 3 to 7 training data sets were generated with different parameters. Starting from the initial model (a set of weights of the neural network) that was trained with the data that mimics the real experimental data, we trained models with these modified training data sets. Once training was completed, we tested each model with the real data of experiment #1. The best model inherits its weights to the next generation model and new training data sets were generated by increasing the occurrences of the poorly characterized events. For examples, if a model predicts a new fibril as growth of a nearby existing fibril, we increased the number of newly appearing fibrils in the new training. If a model predicts a fast-growing event as the appearance of a new fibril, we increased the growth speed acceleration. This process was repeated until the prediction is indistinguishable from the result by human inspection. The neural network was trained using TensorFlow 1.14 with a Tesla P100 GPU of NIH HPC Biowulf cluster.

##### Hyperparameters

An Adam optimizer^8^ with a learning rate of 0.001 was used with dice loss metric.

#### Determination of fluorescence lifetime of each fibril

The fluorescence lifetime of each fibril was calculated using the mean delay time of the photons contained in the image pixels of a fibril corrected for the mean delay time of background photons and the offset by the instrument response function (IRF) of the detector^9^. We determined the average lifetime of background photons by averaging lifetimes of background pixels, the intensity of which is lower than 90 % of the average count rate of the predicted background image.

#### Estimation of number of strands in a fibril

Solid state NMR structures have revealed the polymorphism of Aβ fibrils, which consist of different number of strands with distinct structures. We estimated the number of strands of each fibril using the length from the image and the number of monomers comprising the fibril.

To determine the length of a fibril, the individual fibril image was rotated to make the fibril axis approximately parallel to the *x*-axis using a linear fitting. Next, pixels were segmented to have a length along the *x*-axis smaller than 10 pixels for a segment. Individual segments were fitted to a 3^rd^ order polynomial from the left to the right with restricting the ends of two consecutive segments are continuously connected. Fibrils shorter than 500 nm (~ twice the size of PSF) were excluded from the further analyses that use length information.

The number of monomers in a fibril (*N*) was calculated by comparing the number of photons and fluorescence lifetime of the fibril (*N*_p,fibril_ and τ_fibril_) with those of the monomer (*N*_p,monomer_ and τ_monomer_) measured at the same illumination intensity to account for the reduced intensity of Alexa 594 in fibrils due to fluorescence quenching as

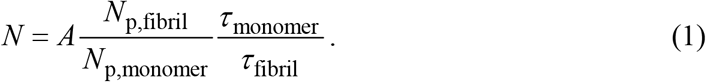

Here, a conversion factor *A* (= 2.7) is used to make the photon densities of different fibril groups are integer multiples of a common photon density because the number of fibril should be an integer. However, small oligomer signals with only a few photons often lead to unreasonable lifetimes values after background correction (shorter than 0 ns or longer than the unquenched monomer lifetime). Therefore, we set the maximum and the minimum value of the lifetimes for the quenching correction in equation (1). The minimum lifetime for the quenching correction is 0.15 ns, which is the 10^th^ percentile of an exponential distribution with the shortest lifetime (1.4 ns) of fibrils with more than 5,000 photons which show strong quenching in the late phase of the experiments. The maximum lifetime for the quenching correction is 3.67 ns, the unquenched lifetime of the monomer.

In the calibration experiment, a direct measurement of the number of photons from the monomer after excitation at 485 nm is not possible due to the very low fluorescence intensity from the monomer at low illumination intensity for fibril imaging. Therefore, we used pulse-interleaved excitation using two picosecond-pulsed lasers (485 nm, LDH-D-C-485 and 595 nm, LDH-D-TA-595, PicoQuant). Monomers can be easily identified in an image collected using 595 nm excitation. The pixels comprising each monomer image are saved and used for the calculation of photons emitted by 485 nm excitation after subtraction of the background. The average number of photons emitted from the monomer, *N*_p,monomer_ = 1.60 and the lifetime of Alexa 594 attached to the monomer is τ_monomer_ = 3.67 ns.

#### Clustering of two-dimensional plot of lifetime and photon density

Fibrils from five experiments were clustered into three groups using Gaussian mixture models^10^ (Fig. 4b). However, we observed that photon density slightly fluctuates in different experiments. Therefore, we normalized photon counts using group 1 (the longest lifetime group) which shows the smallest overlap with other groups in the 2D plot. After normalization, we calculated the average photon density of each group and conversion factors to convert the number of photons to the number of monomers. Since the ratios of the photon densities of Group 1, 2, and 3 are close to 2:4:3, we assumed that the number of stands of the three groups are 2, 4, and 3, respectively, in the calculation of the conversion factors. The average of the conversion factors of the three groups was used in further analyses (*A* in equation (1)). Short fibrils without length information were clustered using their lifetime distances from the average lifetimes of the fibril groups (group 1: τ > 2.04 ns, group 2: 1.35 ns < τ ≤ 2.04 ns, group 3: τ ≤ 1.35 ns).

#### Fibril growth analysis

In the analysis of individual fibril growth, we removed the frames of a fibril containing less than 10 photons which can result from the background fluctuation. When a fibril grows and reaches the boundary region (three pixels from the edge of a 10 × 10 μm^2^ image), we also removed the rest of image frames for that fibril. After deletion, we selected the longest continuous frame sequence for the movie of each fibril.

In the growing and non-growing fibril analysis, to minimize the possibility of including stochastically slowly-growing fibrils in the non-growing group, the 5^th^ percentile of the average growth speed of length-characterizable fibrils was used for the separation criterion for growing and non-growing fibrils (128 monomers/hour or 63.4 photons/h).

For the lifetime histograms, lifetimes were calculated from more than 200 photons.

---

**Supplementary Fig. 1.**
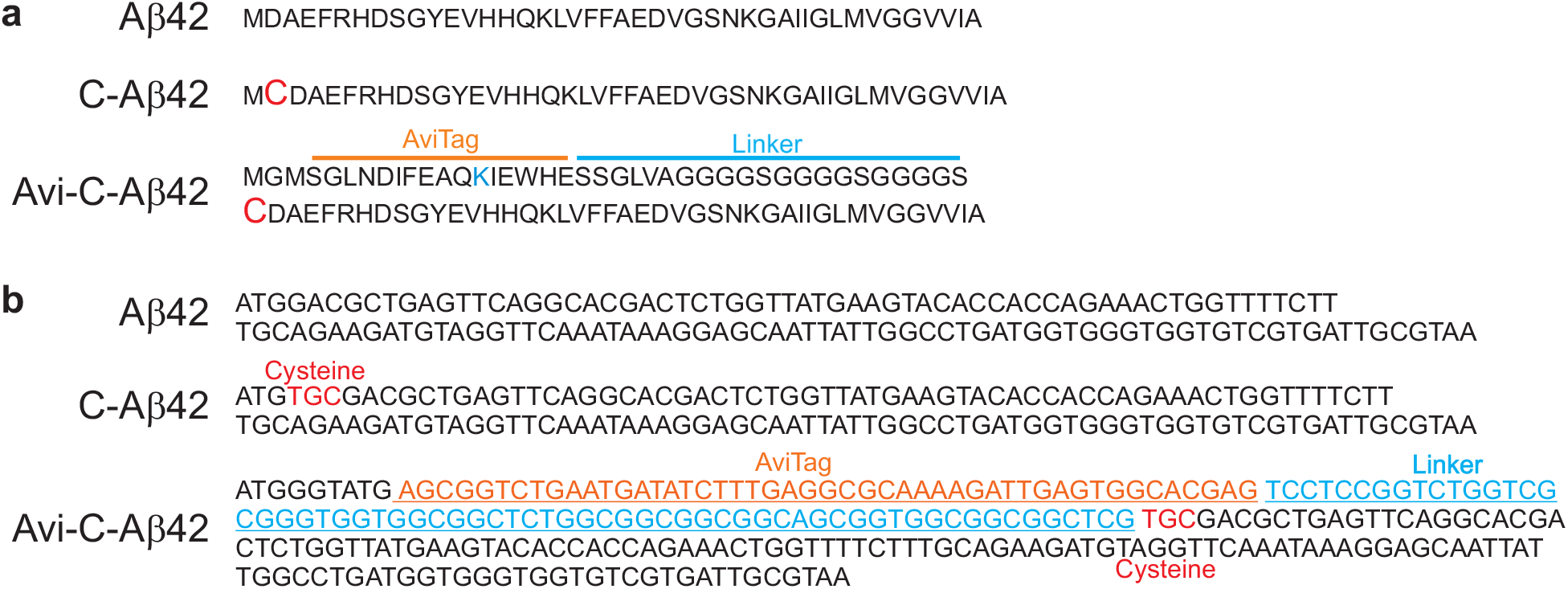
Amino acid and DNA sequences of Aβ42 constructs. **a**, Amino acid sequence. A cysteine residue (magenta C) is appended to the N-terminus for C-Aβ42 and Avi-C-Aβ42. Biotin is attached to the lysine residue (blue K) in the AviTag sequence in the Avi-C-Aβ42 construct, which is separated from Aβ42 sequence by a flexible linker. **b**, DNA sequences.

**Supplementary Fig. 2.**
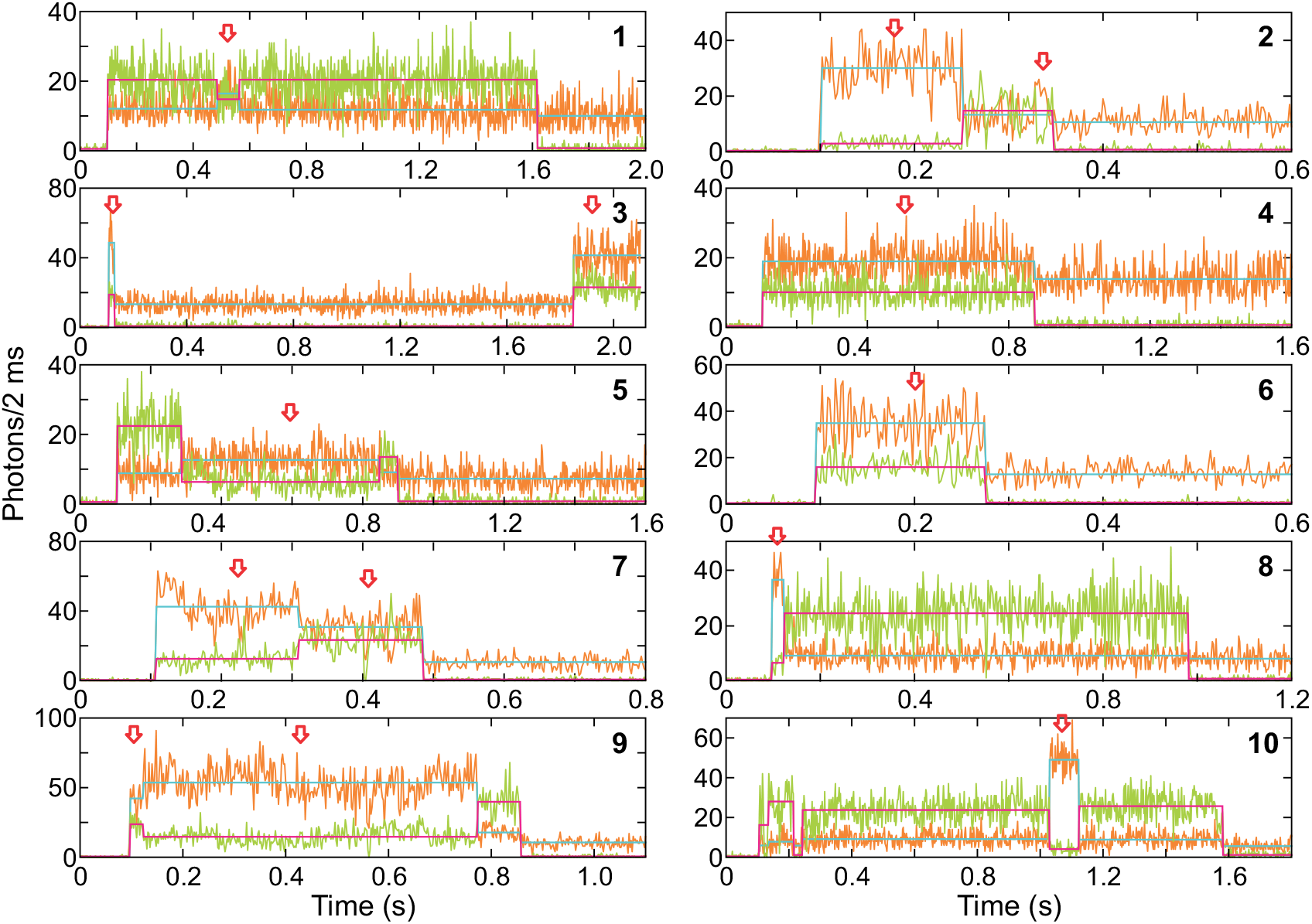
Dimer trajectories of Aβ42. The donor and acceptor fluorescence trajectories (2 ms bin time) of Aβ dimers. Open red arrows indicate dimer segments. Magenta and cyan lines indicate the mean photon count rates of donor and acceptor segments, respectively. Trajectory 4, 6, and 8 show a constant level of fluorescence intensity from the dimer that appears from the beginning of the trajectory (There is a delay of ~ 100 ms between the beginning of data collection and the beginning of illumination to prevent photobleaching before the data collection). Trajectory 2 and 3 begin with the dimer state and exhibit two segments of the dimer state separated by donor-only segments. The appearance of the donor-only segment in the middle of two dimer segments may result from acceptor blinking rather than the dissociation of the dimer and re-association due to the slow dimerization kinetics. Trajectory 7 and 9 begin with the dimer state, but a transition occurs to another dimer state. This can be conformational changes of the dimer or may result from the formation or dissociation of oligomers larger than the dimer. For example, the fluorescence intensity changes in Trajectory 7 looks like double acceptor photobleaching, indicating this can be a trimer. Seven trajectories begin with the dimer state, which were used for the calculation of the dimer dissociation constant. Trajectory 1, 5, and 10 begin with the monomer and dimerization occurs, which is followed by dissociation or photobleaching of the acceptor.

**Supplementary Fig. 3.**
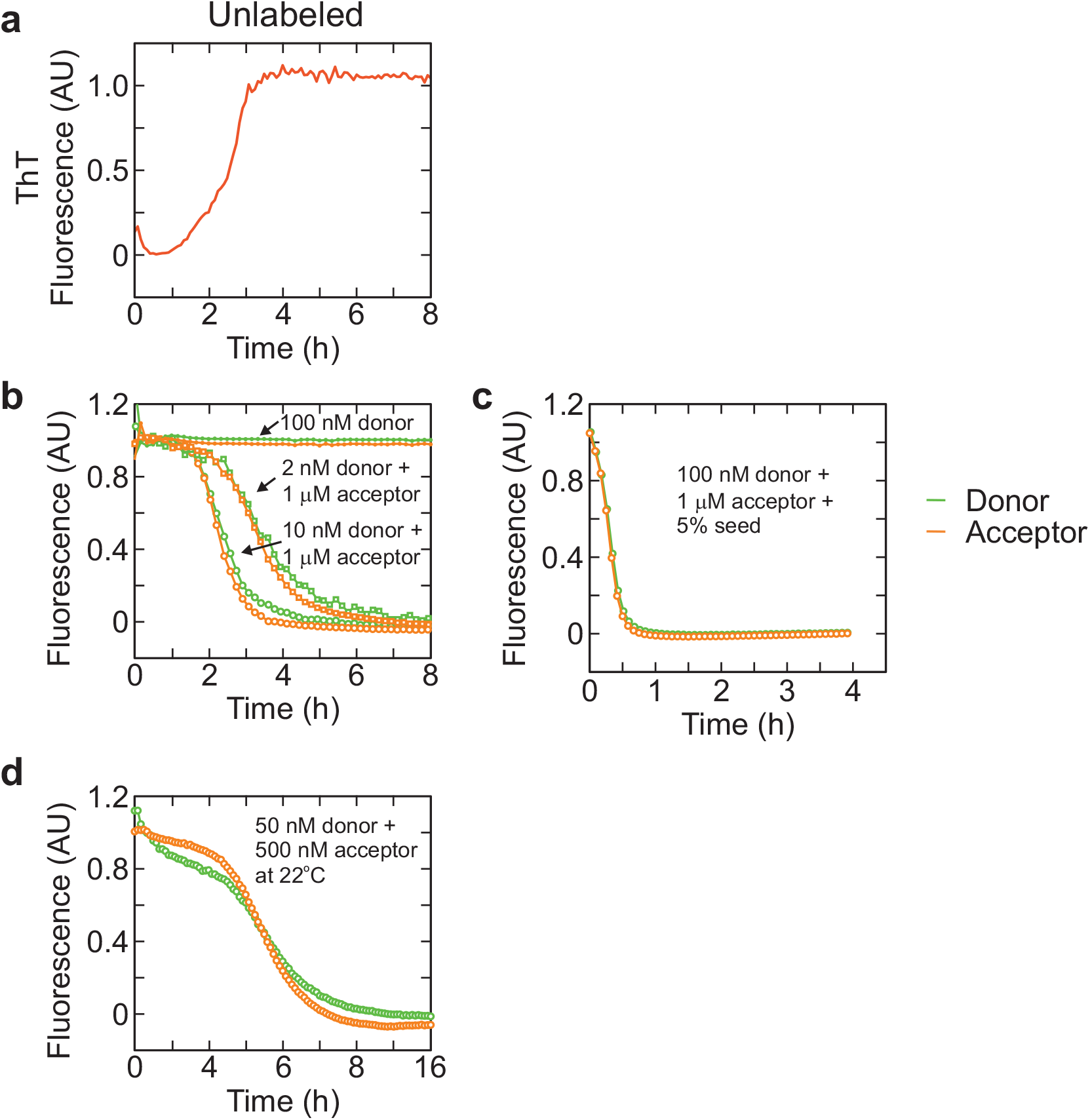
Aggregation of Aβ42. **a**, Aggregation of unlabeled Aβ42 (2 μM) at 37°C monitored by Thioflavin T (ThT) fluorescence. **b**, Co-aggregation of 1 μM of acceptor-labeled Aβ42 with 2 or 50 nM of donor-labeled Aβ42 at 37°C. 100 nM of donor-labeled Aβ42 did not aggregate for 8 hours. **c**, 5% addition of sonicated aggregate of dye-labeled Aβ42 eliminated the lag phase prior to aggregation (37°C). **d**, Aggregation of Aβ42 at a lower concentration (500 nM) and room temperature (22°C), which is the condition for FLIM imaging of Aβ42 aggregation.

**Supplementary Fig. 4.**
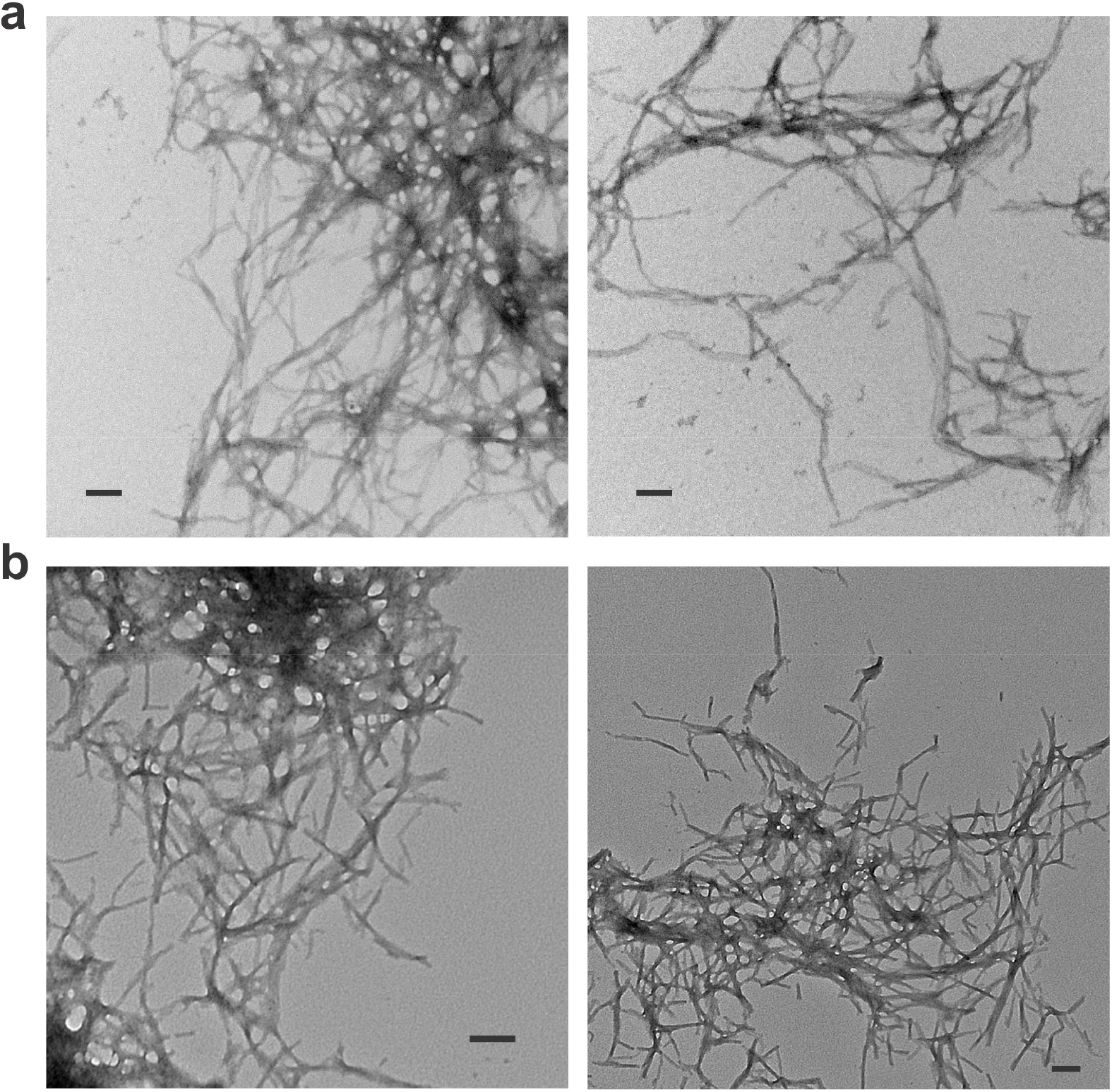
Electron microscope images of fibrils. **a**, Fibrils formed from 5 μM unlabeled Aβ42 at 37°C overnight. **b**, Fibrils formed from 5 μM Alexa 594-labeled Aβ42 at 37°C overnight. Scale bars, 100 nm.

**Supplementary Fig. 5.**
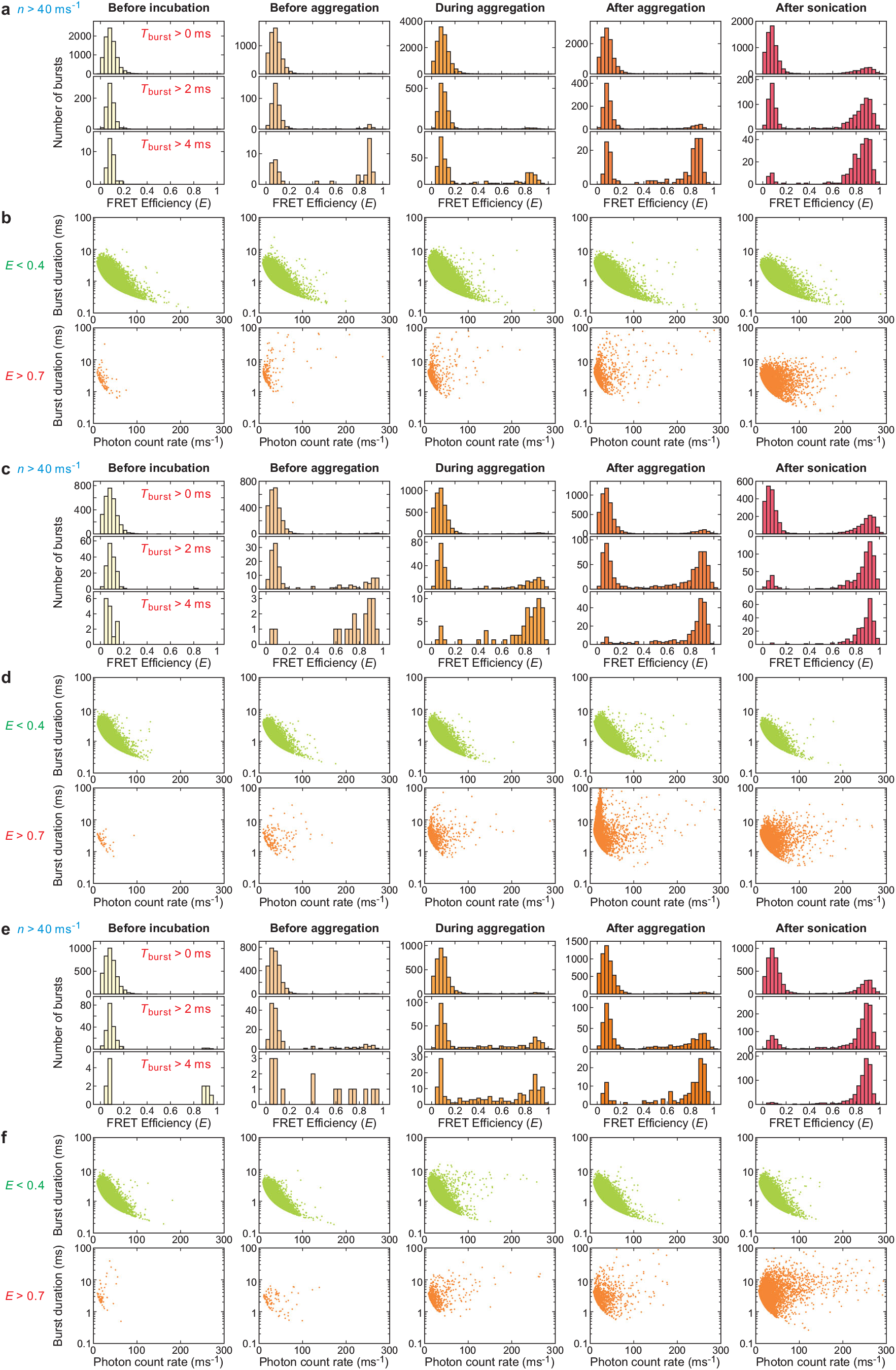
Single-molecule free-diffusion experiment of Avi-D-Aβ42 and A-Aβ42 mixture with 1:10 ratio. 100 nM Avi-D-Aβ42 was mixed with 1 μM A-Aβ42 and incubated at 37°C in a plate reader. A small amount of solution was collected from the plate reader for the free-diffusion experiment at four time points, before incubation, before aggregation, during aggregation, and after aggregation determined from the fluorescence intensity profile (see Fig. 2b). After aggregation is finished, the sample was sonicated and diluted for the free-diffusion experiment. (**a, c, e**) FRET efficiency histograms of fluorescence bursts with the average count rate greater than 40 ms-1 and with three different criteria for the burst duration (*T*_burst_). (**b, d, f**) Two-dimensional plots of the photon count rate and duration of individual bursts with *E* < 0.4 (i.e. donor-only monomer) and *E* > 0.7 (oligomers and fibrils). The experiment was performed in triplicate: **a, b**, blue, **c, d**, red, and **e, f**, yellow traces in Fig. 2b. The protein concentrations in terms of the monomer for the five experiments (from the left to the right) are **a, b**, 200 pM, 200 pM, 200 pM, 200 pM, and 200 pM, **c, d**, 200 pM, 200 pM, 500 pM, 1 nM, and 1 nM, and **e, f**, 200 pM, 200 pM, 500 pM, 1 nM, and 1 nM.

**Supplementary Fig. 6.**
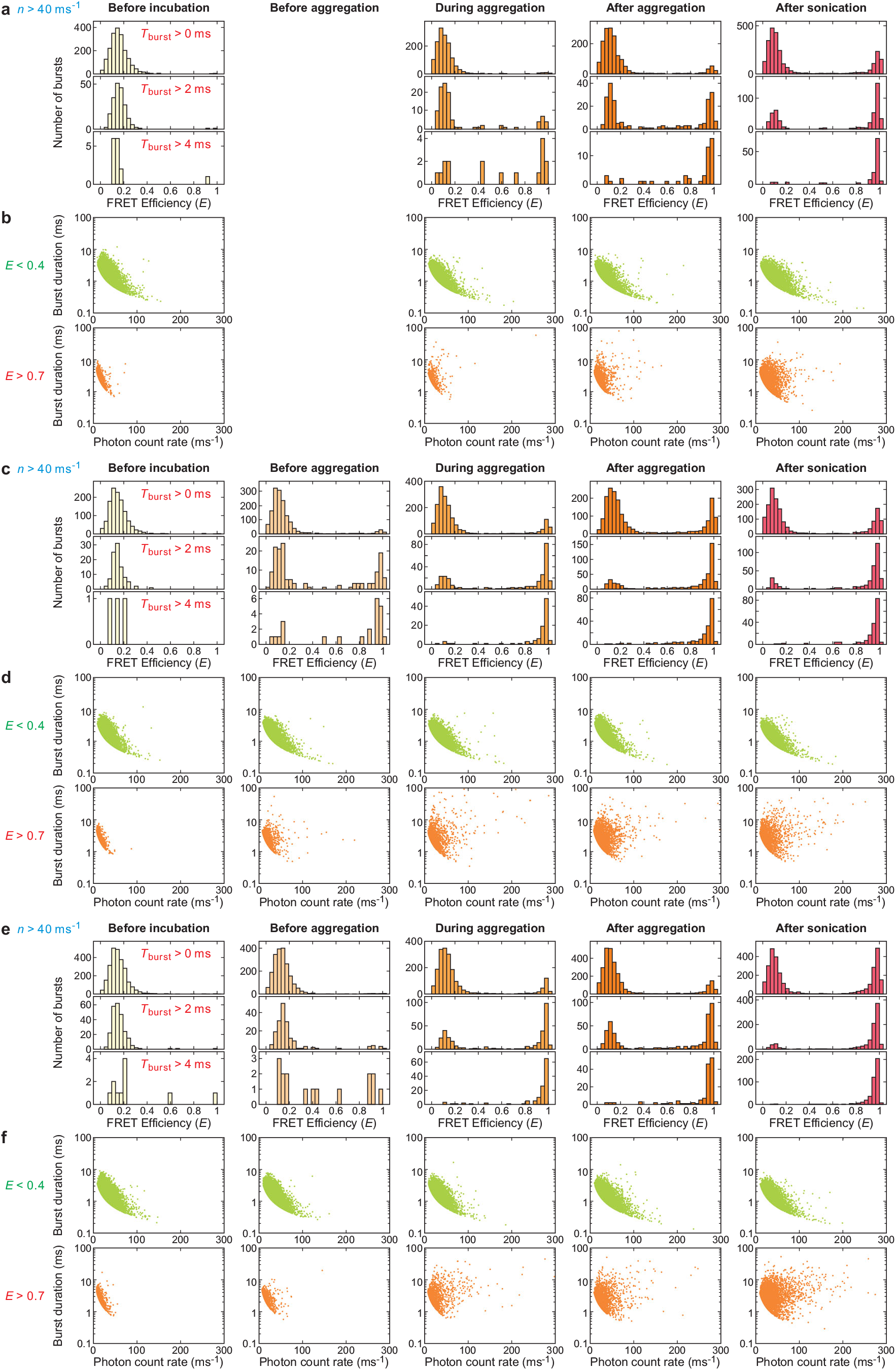
Single-molecule free-diffusion experiment of Avi-D-Aβ42 and A-Aβ42 mixture with 1:100 ratio. 10 nM Avi-D-Aβ42 was mixed with 1 μM A-Aβ42 and incubated at 37°C in a plate reader. A small amount of solution was collected from the plate reader for the free-diffusion experiment at four time points, before incubation, before aggregation, during aggregation, and after aggregation determined from the fluorescence intensity profile (see Fig. 2b). After aggregation is finished, the sample was sonicated and diluted for the free-diffusion experiment. (**a, c, e**) FRET efficiency histograms of fluorescence bursts with the average count rate greater than 40 ms-1 and with three different criteria for the burst duration (*T*_burst_). (**b, d, f**) Two-dimensional plots of the photon count rate and duration of individual bursts with *E* < 0.4 (i.e. donor-only monomer) and *E* > 0.7 (oligomers and fibrils). The experiment was performed in triplicate: **a, b**, blue, **c, d**, red, and **e, f**, yellow traces in Fig. 2b. The protein concentrations in terms of the monomer for the five experiments (from the left to the right) are 50 pM, 50 pM, 100 pM, 200 pM, and 100 pM. The data before aggregation in **a** and **b** could not be collected because the aggregation (blue trace in Fig. 2b) happened before 1 hour, the first time point of sample collection.

**Supplementary Fig. 7.**
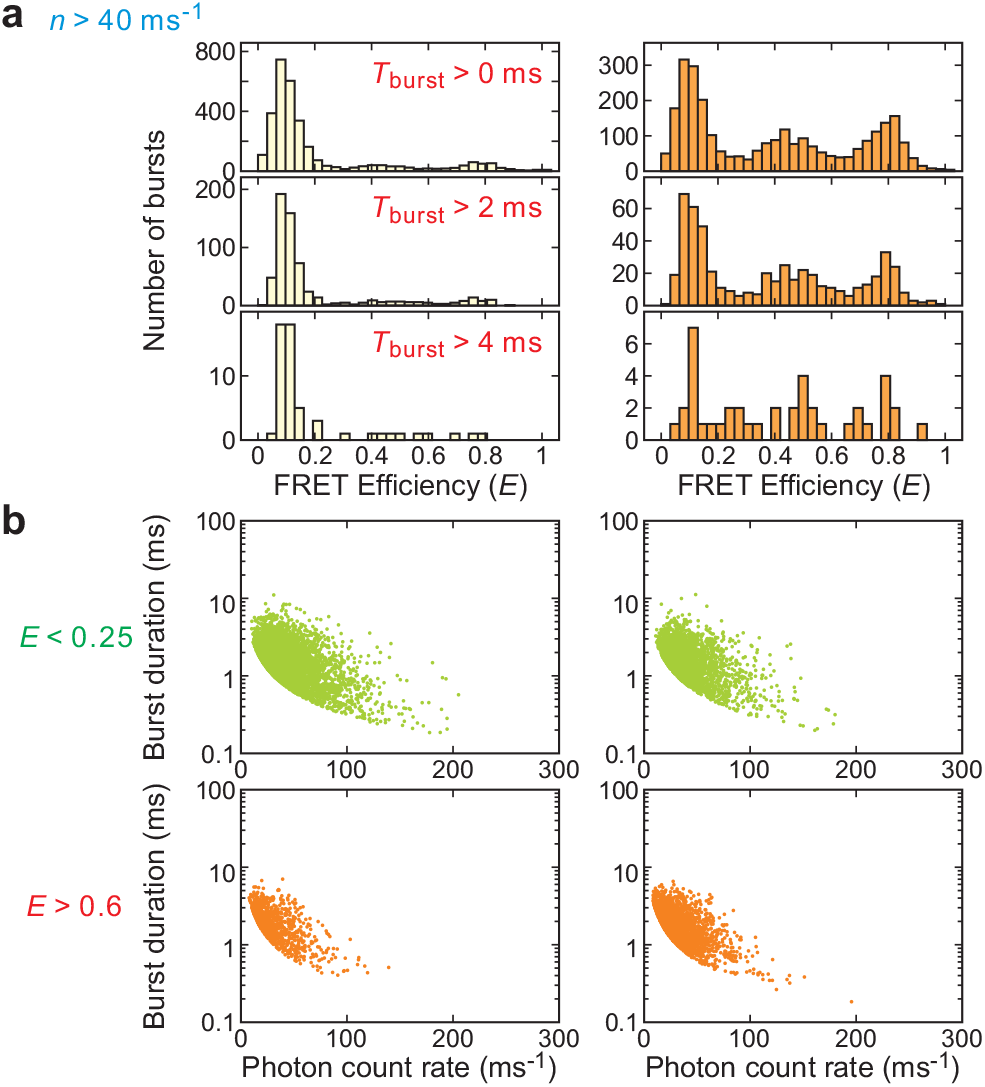
Single-molecule free-diffusion experiment of the tetramerization domain (TD) of p53. 100 nM acceptor-labeled TD at its C-terminus (TD-A) was diluted into 40 pM donor-labeled TD at the N-terminus (Avi-D-TD). The final concentration of TD-A was 5 nM. The data was analyzed for the fluorescence bursts collected within 30 min after mixing (i.e., mostly monomer, left) and between 120 and 150 min after mixing (equilibrium between monomer, dimer, and tetramer, right). **a**, FRET efficiency histograms of fluorescence bursts with average count rate greater than 40 ms-1 and with three different criteria for the burst duration (*T*_burst_). **b**, Two-dimensional plots of the photon count rate and duration of individual bursts with *E* < 0.4 of the data collected within 30 min in **a** (i.e. donor-only monomer) and *E* > 0.6 of the data collected between 120 and 150 min in **a** (i.e., tetramer).

**Supplementary Fig. 8.**
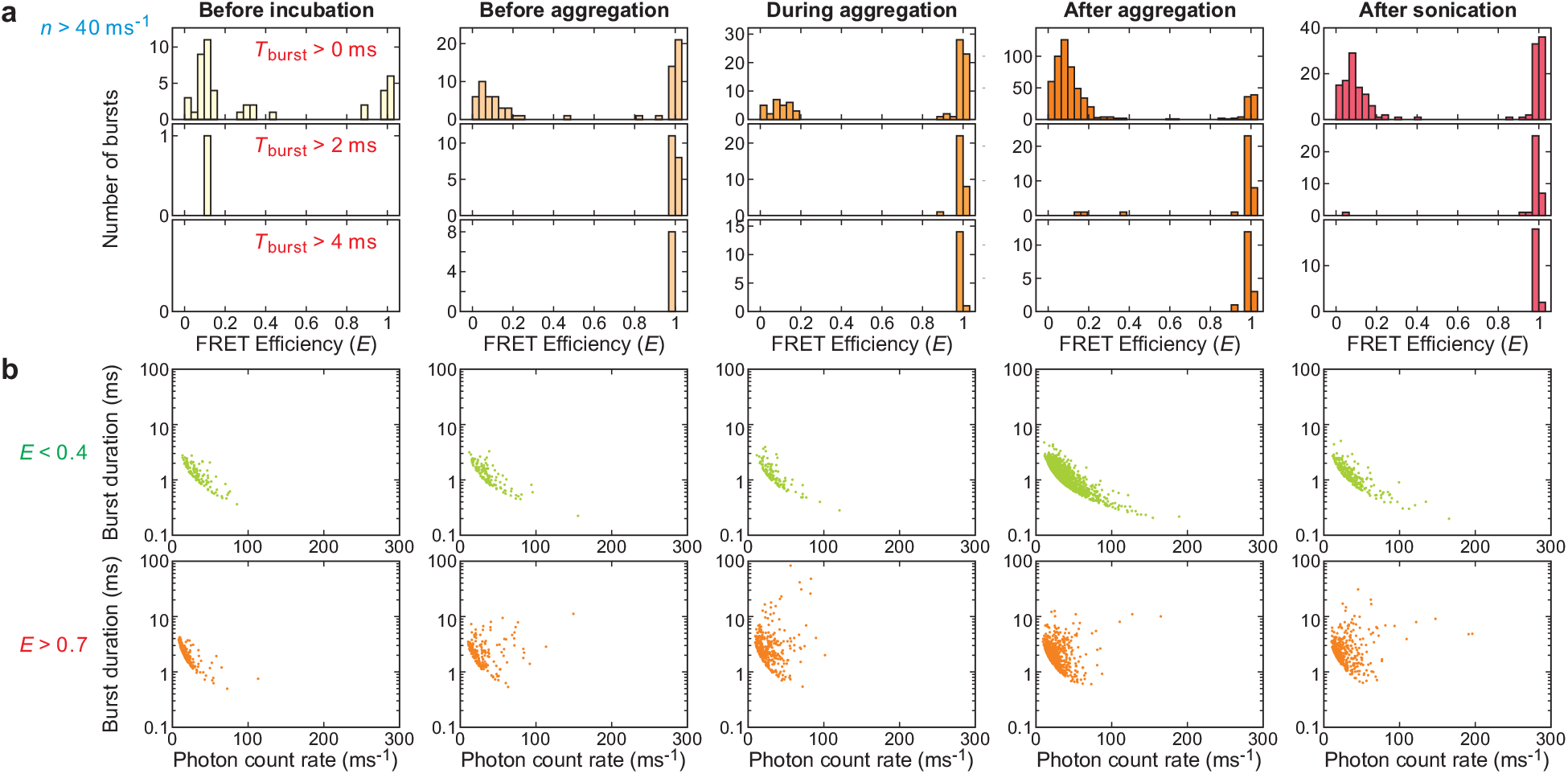
Single-molecule free-diffusion experiment of Alexa 594-labeled Aβ42 (1 μM). The experiment was performed exactly the same as 1:100 mixture experiment in Supplementary Fig. 5 except for the absence of donor-labeled Aβ42. **a**, FRET efficiency histograms of fluorescence bursts with the average count rate greater than 40 ms-1 and with three different criteria for the burst duration (*T*_burst_). **b**, Two-dimensional plots of the photon count rate and duration of individual bursts with *E* < 0.4 and *E* > 0.7. The protein concentrations in terms of the monomer for the five experiments (from the left to the right) are 50 pM, 50 pM, 100 pM, 200 pM, and 100 pM.

**Supplementary Fig. 9.**
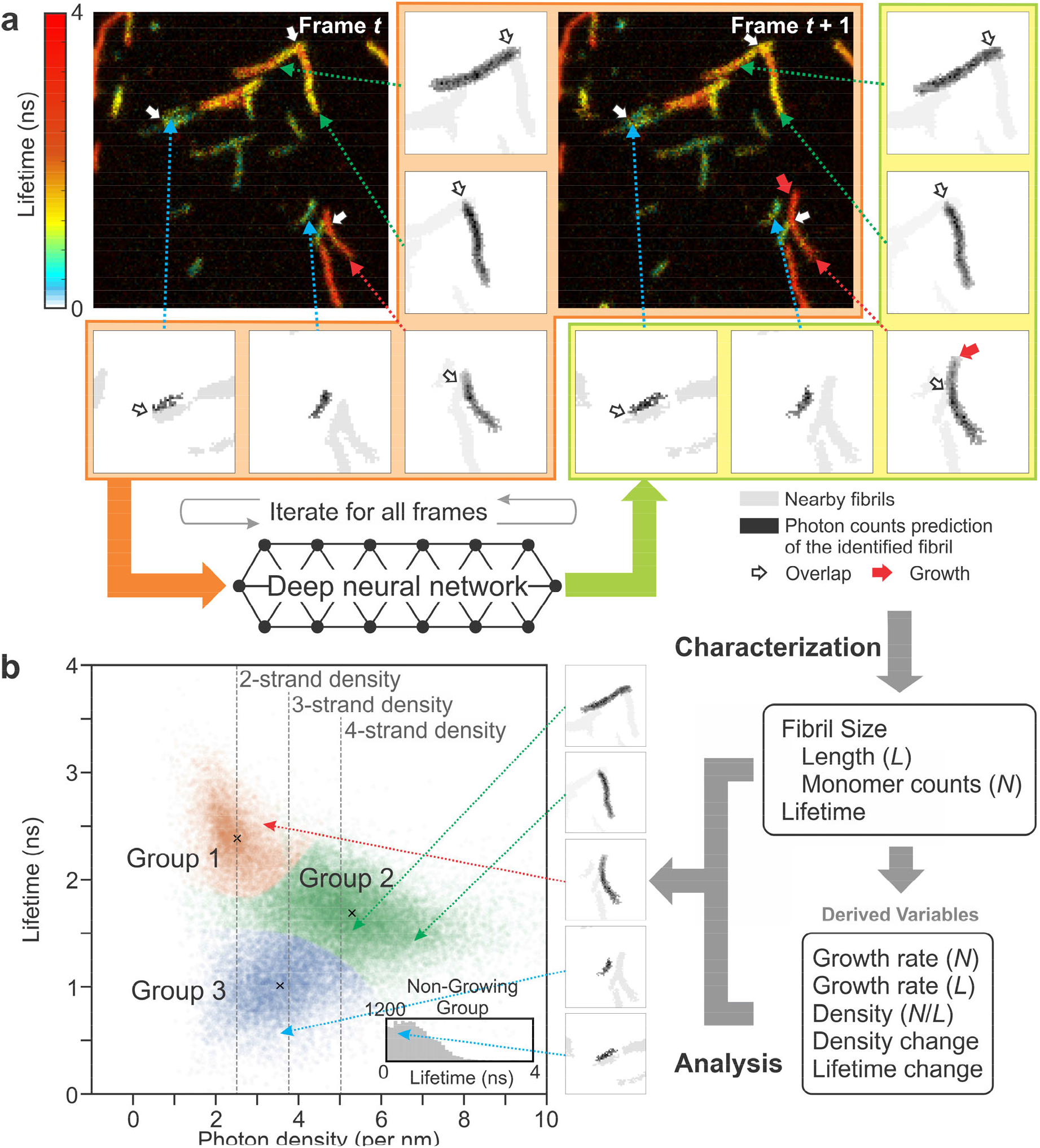
Flowchart of individual fibril analysis. **a**, Identification and separation of individual fibrils by using a deep neural network. Individual fibril images (5 images as examples) of frame *t* and the new image of frame *t* + 1 are processed in deep neural network (images inside the orange area), which results in the updated individual fibril images (inside the yellowish green area) and new fibrils (not shown). The procedure is repeated for the entire time series of images of the same region (see Supplementary Fig. 10 for more complete description of deep learning). Five individual fibrils with different characteristics are shown in the sub-regions on the right and bottom side of the image as examples. Each sub-region consists of only one fibril (black), but nearby fibrils (grey) are also shown for comparison. White arrows indicate the regions where fibrils overlap. Red arrow in the individual image on the lower right corner indicates the growth of a fibril. **b**, (Right) After separation, fibrils in each image are characterized by various properties: length (*L*) measured on the fibril image, fluorescence lifetime, and size in terms of the number of monomer (*N*), which is calculated using the fluorescence intensity and lifetime. Using these parameters from individual images, other time-dependent variables are extracted: growth rate in terms of the length and size (number of monomer) and changes of the monomer density and lifetime of fibrils. (Left) Using the variables from these characterizations, fluorescence lifetime of Alexa 594 attached to Aβ42 vs. photon density is constructed, which is also shown in Fig. 4b. Arrows indicate the locations of fibrils in the 2D plot.

**Supplementary Fig. 10.**
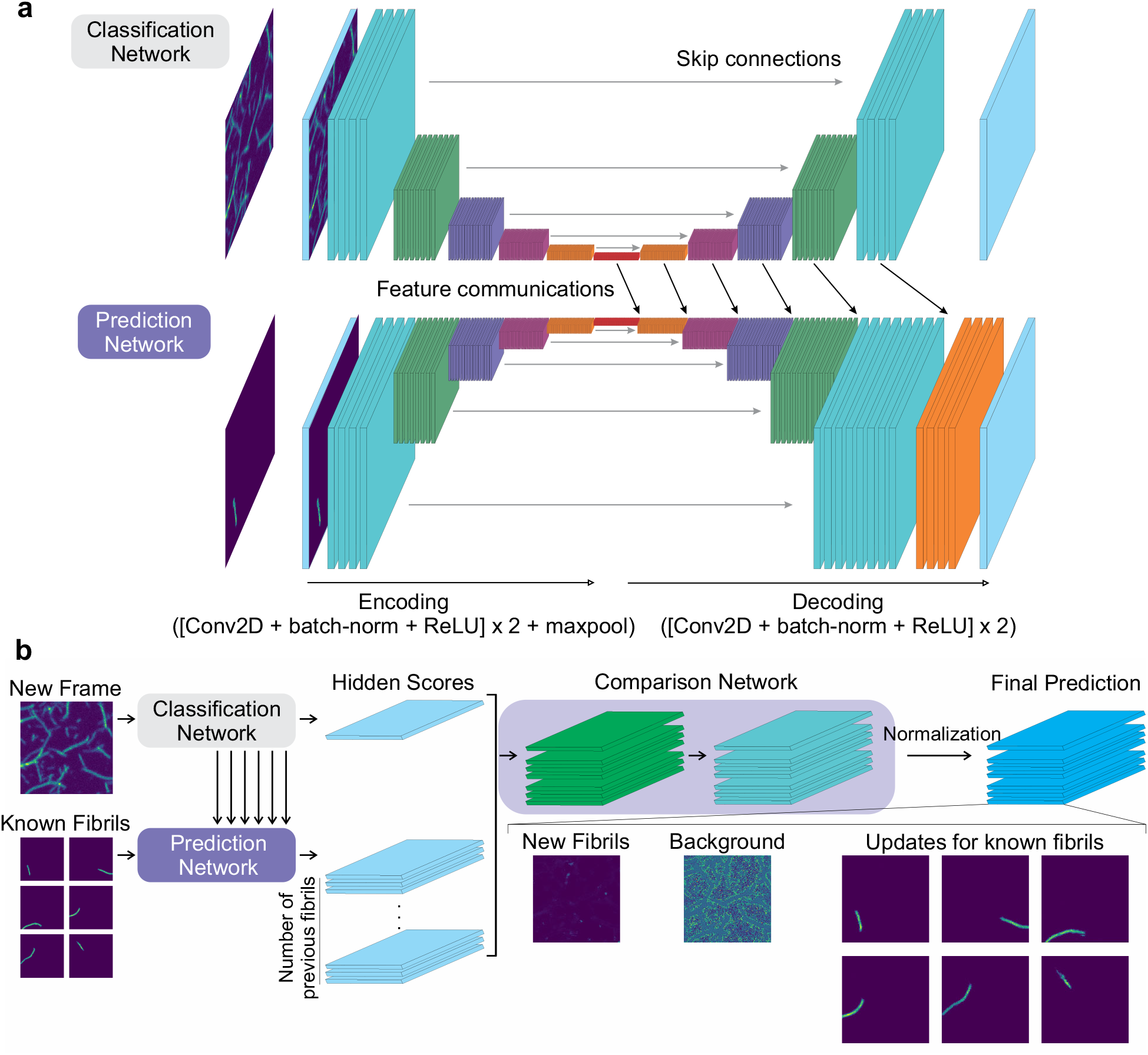
Deep neural network architecture. **a**, The classification network and the prediction network. Each network takes an image as an input. The input of the classification network is a new frame image. The background prediction network uses a background prediction image of the previous frame as an input, and the growth prediction network uses prediction images of individual fibrils of the previous frame (known fibrils) as an input. The prediction network connects the hidden features of the classification network to generate predictions (feature communications). **b**, The output of the classification network and the prediction networks are compared to generate the final prediction. The comparison network has twice applications of a convolution, a batch-normalization, and an ReLU activation layers followed by a convolution and a PReLU activation layer. The output of PReLU activation is normalized pixel-by-pixel to make the number of photons in each pixel of the summed output image equal to that of the original input image of the new frame. This results in an image of new fibrils, a background image, and the updated images of known fibrils from the previous frame (see Supplementary Fig. 7a).

**Supplementary Fig. 11.**
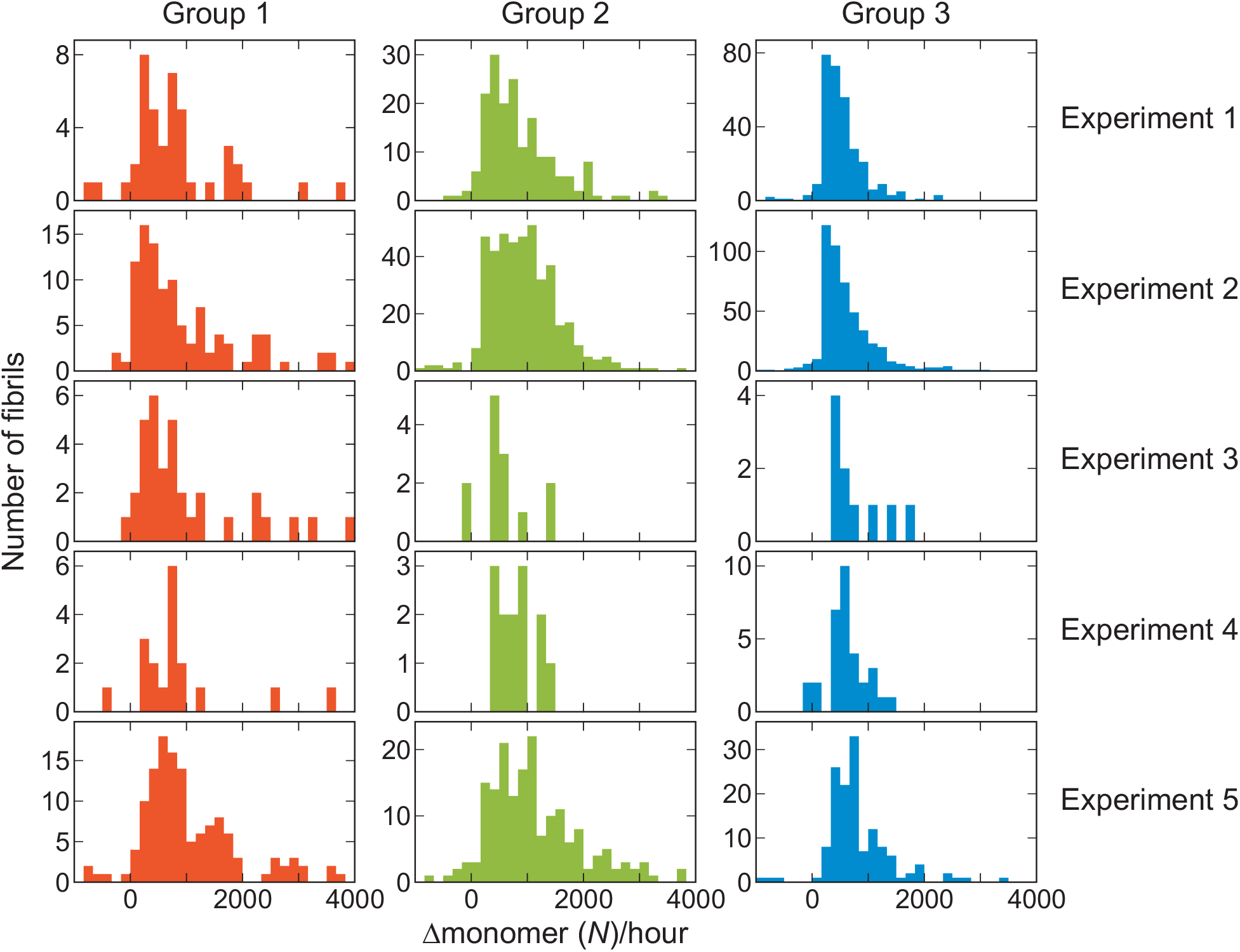
Growth rate of long fibrils of the three fibril groups in individual experiments.

**Supplementary Fig. 12.**
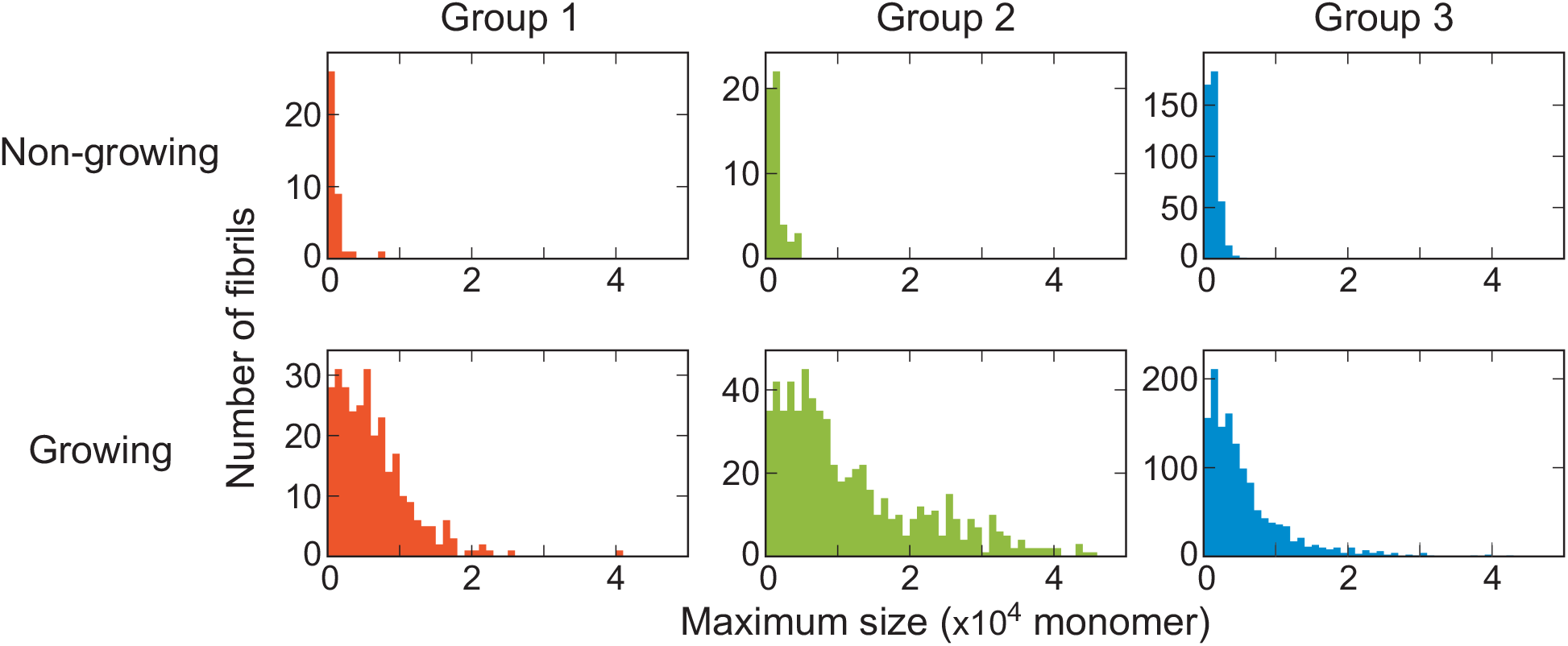
Distribution of the maximum size of fibrils. The maximum size observed for each individual non-growing (upper) and growing (lower) fibrils of the three fibril groups.

**Supplementary Fig. 13.**
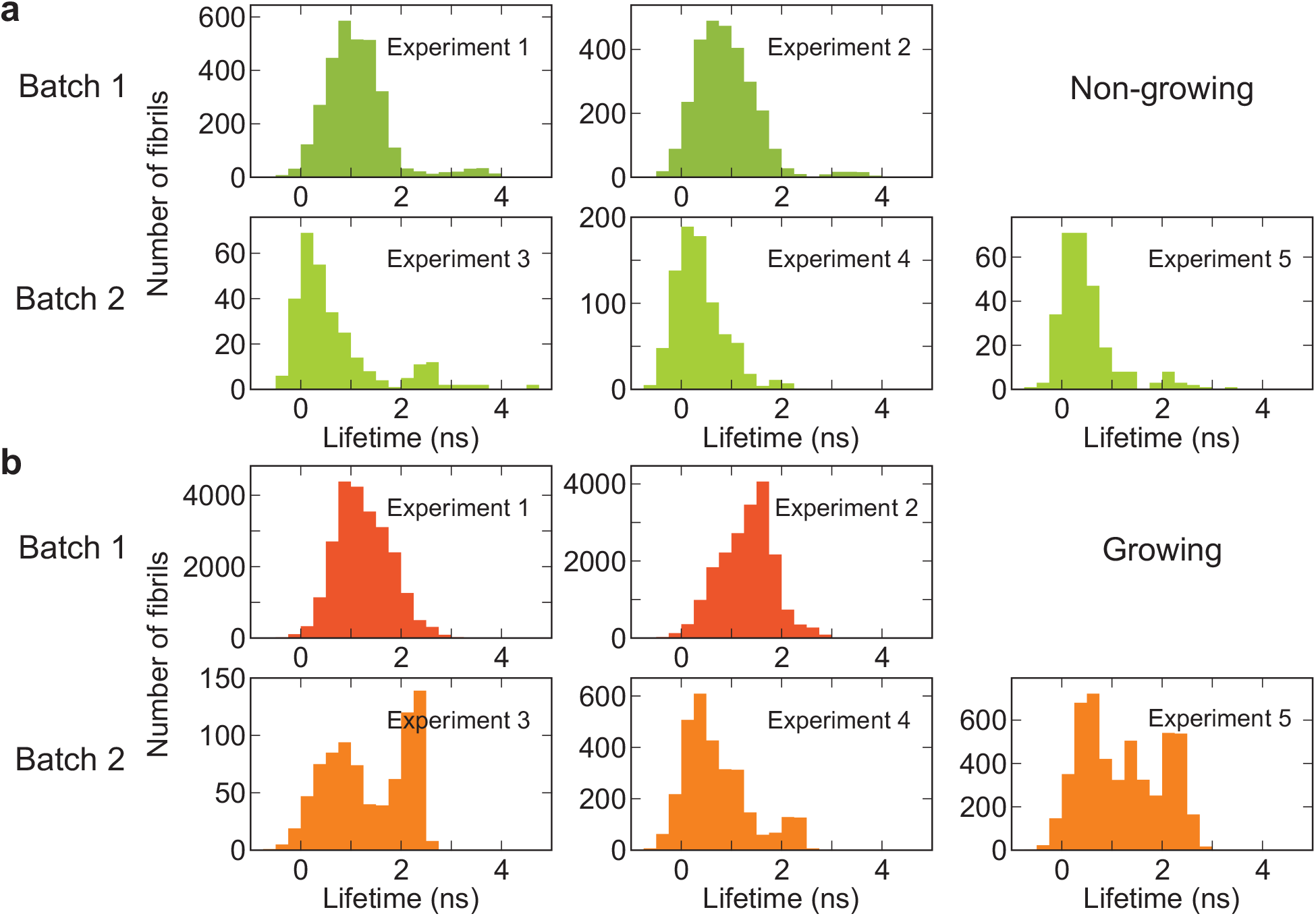
Fluorescence lifetime distributions of five individual experiments. **a**, Non-growing fibrils. **b**, Growing fibrils.

**Supplementary Fig. 14.**
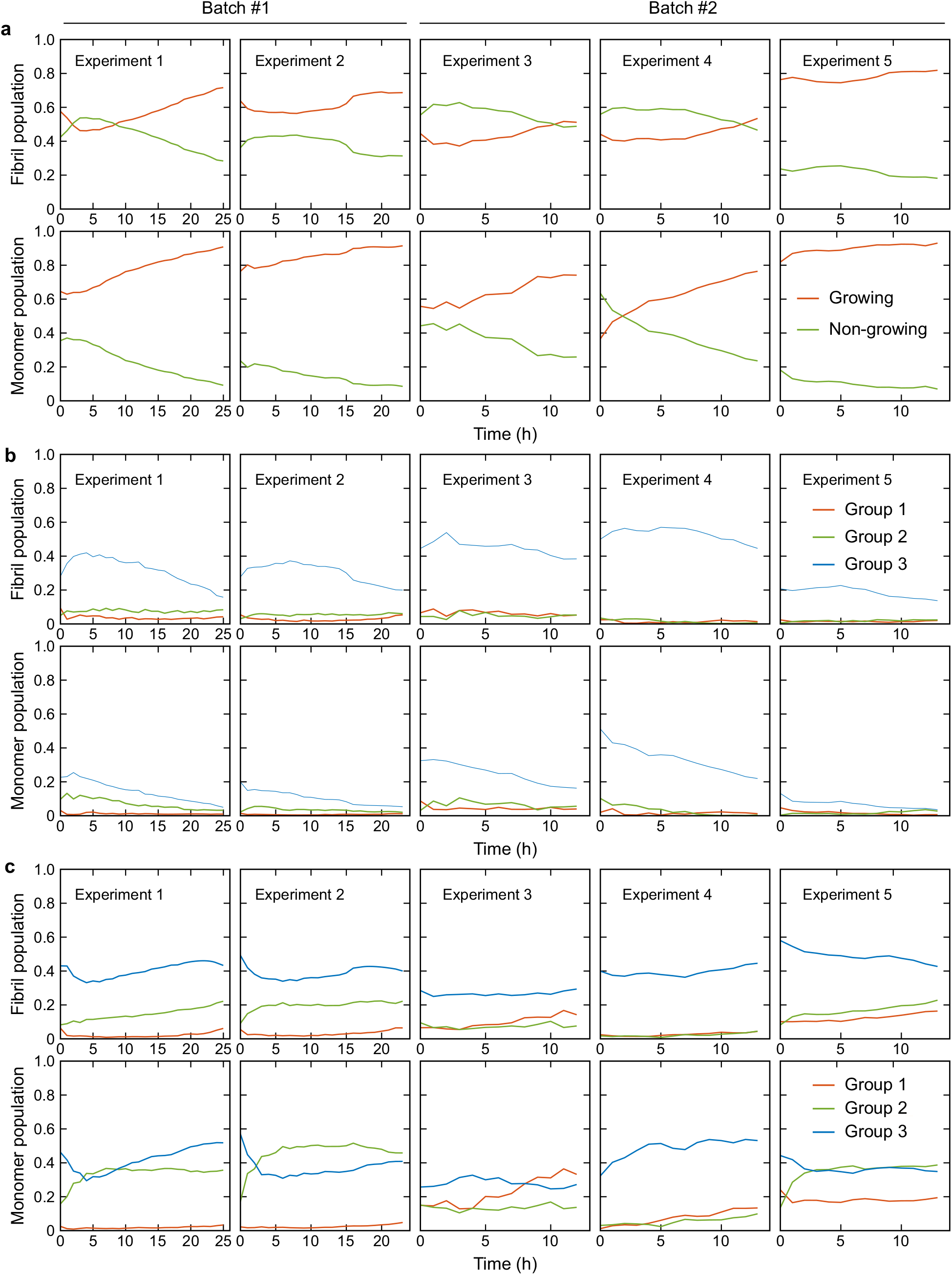
Time-dependent changes of the fractions of monomers in various groups of fibrils from five different experiments. Upper panels show the fraction of number of fibrils and lower panels show fraction of monomers that belong to each fibril category. **a**, Non-growing (green) and growing (red) fibrils. **b**, Three fibril groups of non-growing fibrils. **c**, Three fibril groups of growing fibrils. The fractions are normalized to the total number of fibrils at each time point.

